# Genetically Defined Subtypes of Somatostatin-Containing Cortical Interneurons

**DOI:** 10.1101/2023.02.02.526850

**Authors:** Rachel E. Hostetler, Hang Hu, Ariel Agmon

## Abstract

Inhibitory interneurons play a crucial role in proper development and function of the mammalian cerebral cortex. Of the different inhibitory subclasses, dendritic-targeting, somatostatin-containing (SOM) interneurons may be the most diverse. Earlier studies used transgenic mouse lines to identify and characterize subtypes of SOM interneurons by morphological, electrophysiological and neurochemical properties. More recently, large-scale studies classified SOM interneurons into 13 morpho-electro-transcriptomic (MET) types. It remains unclear, however, how these various classification schemes relate to each other, and experimental access to MET types has been limited by the scarcity of type-specific mouse driver lines. To begin to address these issues we crossed Flp and Cre driver mouse lines and a dual-color combinatorial reporter, allowing experimental access to genetically defined SOM subsets. Brains from adult mice of both sexes were retrogradely dye-labeled from the pial surface to identify layer 1-projecting neurons, and immunostained against several marker proteins, allowing correlation of genetic label, axonal target and marker protein expression in the same neurons. Using whole-cell recordings ex-vivo, we compared electrophysiological properties between intersectional and transgenic SOM subsets. We identified two layer 1-targeting intersectional subsets with non-overlapping marker protein expression and electrophysiological properties which, together with a previously characterized layer 4-targeting subtype, account for about half of all layer 5 SOM cells and >40% of all SOM cells, and appear to map onto 5 of the 13 MET types. Genetic access to these subtypes will allow researchers to determine their synaptic inputs and outputs and uncover their roles in cortical computations and animal behavior.

**SIGNIFICANCE STATEMENT:** Inhibitory neurons are critically important for proper development and function of the cerebral cortex. Although a minority population, they are highly diverse, which poses a major challenge to investigating their contributions to cortical computations and animal and human behavior. As a step towards understanding this diversity we crossed genetically modified mouse lines to allow detailed examination of genetically-defined groups of the most diverse inhibitory subtype, somatostatin-containing interneurons. We identified and characterized three somatostatin subtypes in the deep cortical layers with distinct combinations of anatomical, neurochemical and electrophysiological properties. Future studies could now use these genetic tools to examine how these different subtypes are integrated into the cortical circuit and what roles they play during sensory, cognitive or motor behavior.

## INTRODUCTION

In humans as in other mammals, the neocortex is where incoming sensory information relayed from the sensory periphery via the thalamus is processed and perceived, where decisions about appropriate motor responses are made, and where such motor actions are planned and controlled. While the majority of cortical neurons are excitatory, inhibitory interneurons are crucial for proper neocortical development and function (Isaacson and Scanziani, 2011; Le Magueresse and Monyer, 2013; Kepecs and Fishell, 2014). Inhibitory cortical interneurons fall into four main non-overlapping subclasses characterized by expression of the proteins parvalbumin (PV), somatostatin or vasointestinal protein (VIP) or of the gene Inhibitor of DNA binding 2 (*Id2*) (Rudy et al., 2011; Tremblay et al., 2016; Schuman et al., 2019; Machold et al., 2022). In contrast to soma-, proximal dendrites- or axon initial segment-targeting PV cells, which can powerfully silence the final spike output of a neuron, SOM inhibition is more subtle, as it targets distal dendrites (Freund and Buzsaki, 1996; Kawaguchi and Kubota, 1996; Wang et al., 2004; Bloss et al., 2016), where it can affect integration of synaptic inputs and dampen dendritic excitability (Jadi et al., 2012; Lovett-Barron et al., 2012; Doron et al., 2017). Among functions ascribed to this subclass is generating surround suppression (Adesnik et al., 2012; Kato et al., 2017; Lakunina et al., 2020), promoting long-range coherence and sensory-evoked oscillations (Chen et al., 2017; Veit et al., 2017; Abbas et al., 2018; Hakim et al., 2018), and enabling learning and memory (Makino and Komiyama, 2015; Adler et al., 2019; Artinian et al., 2019; Cummings and Clem, 2020; Dobrzanski et al., 2022). Moreover, dysfunction of SOM interneurons is implicated in a variety of neuropsychiatric and neurodevelopmental disorders (Paluszkiewicz et al., 2011; Fee et al., 2017; Chen et al., 2019; Anderson et al., 2020; Zorrilla de San Martin et al., 2020; Wengert et al., 2021; He et al., 2022).

SOM interneurons may be the most diverse of the four inhibitory subclasses. Previous studies using various transgenic mouse lines identified several genetically-defined SOM subsets with distinct morphological, neurochemical and electrophysiological phenotypes. These included GFP-expressing, layer-4 projecting (non-Martinotti) cells in the X94 mouse line (Ma et al., 2006); GFP-expressing, layer-1 projecting (Martinotti) cells in the GIN and X98 mouse lines (Oliva et al., 2000; Ma et al., 2006); and Cre-expressing Martinotti cells in the Chrna2-Cre, Calb1-Cre and Calb2-Cre mouse lines (He et al., 2016; Hilscher et al., 2017; Nigro et al., 2018). In recent years recognition of SOM diversity was reinforced by large-scale transcriptomic taxonomies. For example, a multimodal classification study (Gouwens et al., 2020) clustered SOM interneurons into 13 MET types, but identified only 15 such types in the other three inhibitory subclasses combined. Unfortunately, much of this diversity remains inaccessible to experimenters for lack of genetic targeting tools. Consequently, the great majority of the many studies to-date examining the roles played by SOM interneurons in sensory processing, motor skill acquisition and associative learning targeted the SOM subclass en-masse, using the Sst-IRES-Cre line (Taniguchi et al., 2011). This line labels all SOM interneurons non-selectively and can also induce some off-target recombination (Hu et al., 2013; Mikulovic et al., 2015; Muller-Komorowska et al., 2020). Clearly, there is major gap between our recognition of the transcriptomic and phenotypic diversity of SOM interneurons and our ability to selectively target specific subtypes for recording, imaging or activity manipulation.

To begin to close this gap, we characterized in detail SOM subsets captured by combinatorial breeding of five driver and two transgenic mouse lines. In adult mice of both sexes, we identified three non-overlapping subsets with distinct axonal targets, marker protein expression and electrophysiological properties, that together account for about half of all SOM interneurons in L5 and for >40% of all SOM interneurons. Our findings call for a renewed effort to generate additional driver lines that can be used combinatorially to provide experimental access to additional SOM subtypes, to characterize their intrinsic properties and synaptic connectivity patterns and uncover their roles in cortical computations and behavior.

## METHODS

### Animals

Mice used for histological experiments were 1-5 (typically 2-3) months old, of both sexes. To label genetically distinct subsets of SOM interneurons we crossed Sst-Flp mice (JAX strain #028579) (He et al., 2016) with one of 4 Cre recombinase-expressing mouse lines: Calb2-Cre (JAX strain #010774) (Taniguchi et al., 2011), Calb1-Cre (JAX strain #028532) (Daigle et al., 2018), Chrna2-Cre (Leao et al., 2012), and Pdyn-Cre (JAX strain #027958) (Krashes et al., 2014). Dual-recombinase progeny were then crossed with the RC::FLTG reporter line (JAX strain #026932) (Plummer et al., 2015), to create triple transgenic mice expressing GFP in Cre+/Flp+ cells and tdTomato in Cre-/Flp+ cells. We also used X94 and X98 mice (JAX strains #006340 and #006334) (Ma et al., 2006) to label previously characterized subsets of L4-projecting and L1-projecting SOM cells, respectively, crossing them with Cre driver lines and the Ai9 tdTomato reporter (JAX strain #007909) (Madisen et al., 2010).

### Retrograde labeling

Mice were deeply anesthetized with isoflurane, placed in a heated stereotactic frame, and injected subcutaneously with local anesthetic (bupivacaine) and analgesic (meloxicam). The skull over the right primary somatosensory (S1, barrel) cortex was exposed and a flap of bone (approx. 2×3 mm) outlined with a ¼ mm drill was removed together with the dura mater. A filter paper circle, pre-saturated with Fast Blue dye solution (FB, Polysciences, Warrington, PA, USA; 1% in distilled water) and allowed to dry, was cut to size, dipped in cortex buffer (composition: 125 mM NaCl, 5 mM KCl, 10 mM glucose, 10 mM HEPES, 2 mM CaCl_2_, 2 mM MgSO_4_), placed over the exposed pial surface and covered with Kwik-Cast silicon sealant (World Precision Instruments). The skin incision was then closed, and mice were allowed to recover. Criteria for successful retrograde labeling are indicated below.

### Histology

One day (24±2 hrs) after surgery, mice were deeply anesthetized with an intraperitoneal injection of Avertin and transcardially perfused with ∼30 ml of room temperature saline followed by 50 ml of room temperature 4% paraformaldehyde (PFA) at 5 ml/min. Brains were removed and post-fixed in 4% PFA at room temperature for 4 hours on a shaker plate, then placed in 30% sucrose in phosphate-buffered saline (PBS) at 4°C on a shaker plate for at least 1-2 days. Brains were sectioned through the barrel cortex on a freezing microtome (-30°C) in the coronal plane at a thickness of 30 μm (60 μm for X94 mice). Every FB-labeled tissue section within barrel cortex was collected. For immunostaining, every 4th section immunostained with each antibody. In total, 4-12 sections were collected per brain, and typically 4 sections were stained with each antibody.

### Immunocytochemistry

Free-floating fixed sections were blocked in 5% goat serum, 0.5% Triton X-100 (TX) in PBS for 1 hour at room temperature, and then incubated with primary antibody in 1.25% goat serum, 0.125% TX in PBS for 48 hours at 4°C. Sections were then washed 3× with PBS and incubated with secondary antibodies in 1% goat serum, 0.1% TX in PBS for 2 hours at room temperature. Finally, sections were washed 3× with PBS and mounted in Vectashield Antifade (Vector Laboratories) or Prolong Diamond (Molecular Probes) mounting medium. Sections stained with mouse primary antibodies were blocked in ReadyProbes “Mouse on Mouse” IgG blocking solution (Thermofisher) for 1 hour before the initial blocking step. Primary antibodies and dilutions used were rabbit anti-calretinin (Swant, 1:2000), mouse anti-calbindin (Swant, 1:500), and rabbit anti-neuropeptide Y (Immunostar, 1:2000). Secondary antibodies used were AlexaFluor 647 anti-rabbit (Thermofisher, 1:1000) and AlexaFluor 647 anti-mouse (Thermofisher, 1:500).

### Confocal imaging and histological analysis

The same sections were used to analyze fluorescent protein expression, FB labeling and immunostaining. Images of stained sections were taken on a Nikon A1R inverted confocal microscope using a 20X, NA 0.75 objective, at a Z-step of 2.5 μm. Lasers of 405, 488, 561, and 640 nm wavelengths were used to excite FB, GFP, TdTomato and AlexaFluor 647, respectively. In each section, a region of interest encompassing the cortical region underlying the FB deposit, typically 700-1000 μm wide, was selected, and all SOM interneurons within it (identified by GFP or tdTomato expression) were digitally marked, using NIS Elements (Nikon), as positive or negative for FB and for the relevant antibody, by visually inspecting the full confocal Z-stack. Cortical layers were determined by cell body shape, size, and density. Only brains with successful retrograde labeling were included in this and subsequent analysis; criteria for successful labeling included strong FB labeling in subplate neurons (as evidence for sufficient incubation time for dye to reach all cortical layers) and largely label-free L4 (as evidence for dye uptake limited to L1). Insufficient labeling typically resulted from incomplete removal of the dura mater or from dislodging of the filter paper during the incubation period, whereas large numbers of FB-labeled cells in L4 typically reflected damage to the pial surface and diffusion of dye to L2/3. Similar selection parameters were used in previous studies using the same retrograde labeling method (Ramos-Moreno and Clasca, 2014).

### Ex-vivo brain slice preparation

Mice (both sexes, typically 1-2 months old) were decapitated under deep isoflurane anesthesia, the brains removed and submerged in ice-cold, sucrose-based artificial cerebrospinal fluid (ACSF) containing (in mM): sucrose 206, NaH_2_PO_4_ 1.25, MgCl_2_.6H_2_O 10, CaCl_2_ 0.25, KCl 2.5, NaHCO_3_ 26 and D-glucose 11, pH 7.4. Thalamocortical brain slices (Agmon and Connors, 1991; Porter et al., 2001) of somatosensory (barrel) cortex, 300-350 μm thick, were cut in same solution using a Leica VT-200 vibratome, and placed in a submersion holding chamber filled with recirculated and oxygenated ACSF (in mM: NaCl 126, KCl 3, NaH_2_PO_4_ 1.25, CaCl_2_ 2, MgSO_4_ 1.3, NaHCO3 26, and D-glucose 20). Slices were incubated for at least 30 minutes at 32°C and then at room temperature until use. For recording, individual slices were transferred to a submersion recording chamber and continuously superfused with 32°C oxygenated ACSF at a rate of 2–3 ml/min.

### Electrophysiological recordings

Recording were done on an upright microscope (FN-1, Nikon) under a 40X water immersion objective. For whole-cell recordings, glass micropipettes (typically 5–8 MΩ in resistance) were filled with an intracellular solution containing (in mM): K-gluconate 134, KCl 3.5, CaCl2 0.1, HEPES 10, EGTA 1.1, Mg-ATP 4, phosphocreatine-Tris 10, and 2 mg/ml biocytin, adjusted to pH 7.25 and 290 mOsm. Labeled neurons were identified visually and on a digital camera (Nikon) by their fluorescence, and targeted for single or dual whole-cell recordings using a MultiClamp 700B amplifier (Molecular Devices, San-Jose, CA, USA). Upon break-in, cells were routinely tested by a standardized family of incrementing 600 ms-long intracellular current steps in both negative and positive directions. In post-hoc analysis, the same records were used to extract multiple electrophysiological parameters for each cell (see below). Data were acquired at a 20 kHz sampling rate using a National Instruments (Austin, TX, USA) ADC board controlled by an in-house acquisition software written in the LabView (National Instruments) environment. Reported intracellular voltages are not corrected for junction potential.

### Post-hoc analysis

A total of 10 electrophysiological parameters were analyzed per cell. Single-spike parameters were measured at rheobase (minimal current evoking an action potential). All current steps were 600 ms long.

Electrophysiological parameters definitions:

**V_rest_:** Resting potential upon break-in, with no holding current applied.

**V_threshold_:** The transmembrane voltage when dv/dt reached 5 V/s.

**Spike height:** Spike peak - V_threshold._

**SWHH** (Spike width at half-height): Spike width measured half-way between V_threshold_ and spike peak.

**AHP**: V_threshold_ - spike trough.

**R_in_**: The slope of the I-V plot, calculated from 4-6 positive and negative subthreshold current steps, at membrane potentials up to ±15 mV from rest.

**Sag:** The slope of the plot of voltage sag (maximum voltage – steady-state voltage) vs. membrane potential, calculated from negative current steps.

**F_max_:** The maximal (initial) firing frequency, computed as the reciprocal of the average of the first 3 ISIs in a spike train elicited by I_max_ (I_max_ being the maximal current step applied before a noticeable reduction in spike height.)

**F_ss_**: The steady-state firing frequency, computed as the reciprocal of the average of the last 5 ISIs in a spike train elicited by I_max_.

**Adaptation Ratio**: F_ss_/F_max_.

### Statistical Analysis

Unless noted otherwise, exact p-values were computed using distribution-free, non-parametric permutation tests, by performing 10,000 random permutations of the data and calculating the fraction of permutations resulting in equal or more extreme values of the relevant statistic (under both tails, except for the F-statistic which is one-sided) (Good, 1999). When no more extreme values were found, this is indicated as p<0.0001. Principal component analysis and discriminant function analysis were computed using custom routines following Manly, 2005. See Ma et al., 2006 for a detailed description of the calculations. All computations were programmed in MathCad; routines are available upon request. All data are reported as mean±SEM unless indicated otherwise.

## RESULTS

### Targeting SOM subsets by intersectional genetics

To develop genetic tools for accessing distinct subtypes of SOM interneurons, we searched for available mouse driver lines in which Cre recombinase was co-expressed with marker genes for identified transcriptomic SOM groups (Tasic et al., 2018). We selected 4 such lines: Calb2-IRES-Cre (Taniguchi et al., 2011), shown to label calretinin (CR)-containing SOM cells (Taniguchi et al., 2011; He et al., 2016; Nigro et al., 2018); Chrna2-Cre (Leao et al., 2012), shown to label hippocampal O-LM interneurons and also a subset of L5 SOM cells expressing the α2 nicotinic receptor subunit (Hilscher et al., 2017); Calb1-IRES2-Cre (Daigle et al., 2018), in which Cre is co-expressed with calbindin (CB), a marker for SOM subsets (Kawaguchi and Kubota, 1997; Ma et al., 2006) and Pdyn-IRES-Cre (Krashes et al., 2014), co-expressing Cre with prodynorphin. We selected the latter line since antibodies to preprodynorphin label a subset of middle-layer SOM cells (Sohn et al., 2014) and we were looking for a line which will target the X94 (non-Martinotti) SOM subtype in these layers (Ma et al., 2006). Since marker protein (and thereby Cre) expression in these lines may not be restricted to SOM interneurons - for example, CR is also expressed by VIP-containing interneurons (Gonchar et al., 2007; Xu et al., 2010), and CB is also weakly expressed by upper layer excitatory cells (van Brederode et al., 1991) - we resorted to an intersectional strategy (He et al., 2016; Nigro et al., 2018; Wu et al., 2022), but chose a novel approach: we crossed each Cre line with the Sst-Flp line and the combinatorial RC::FLTG reporter (Plummer et al., 2015), resulting in triple-transgenic progeny in which the Cre-expressing subset of SOM cells expressed GFP while all other SOM cells (but no other cells) expressed tdTomato (Fig. 1A, Venn diagram at upper right). For convenience, we will refer to the SOM neurons expressing GFP in the 4 intersectional genotypes above as Calb2, Chrna2, Calb1 and Pdyn cells, respectively.

**Figure 1.**
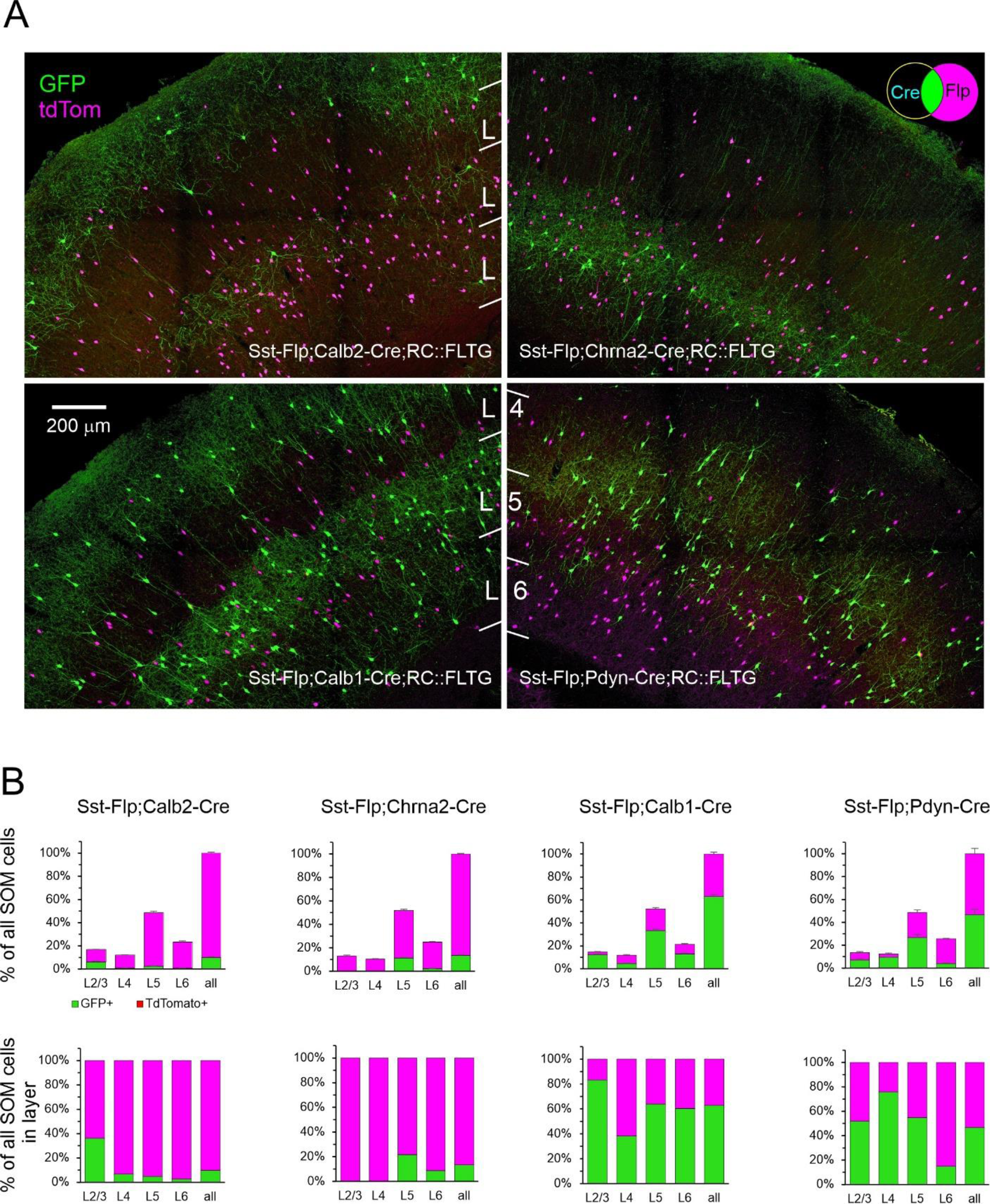
Fluorescent reporter expression in the 4 intersectional genotypes. ***A,*** representative projections of confocal Z-stacks taken with a 20×, 0.75 NA objective at 2.5 μm Z-steps through 30 μm-thick sections. Color channels were adjusted individually. Venn diagram in upper right illustrates the combinatorial logic of the reporter. ***B, upper panels*,** fraction of TdTomato+ and GFP+ cells out of all SOM cells counted in each brain, averaged by genotype. Error bars are SEM. ***Lower panels***, the same data normalized by layer. N=4, 6, 4 and 3 mice for Calb2, Chrna2, Calb1, and Pdyn intersections.

We characterized the 4 intersectional genotypes by imaging and analyzing brains from 17 mice of both sexes, 1-4 months old (except for one animal >5 months), 3-6 mice per genotype. Perfusion-fixed brains were cut into 30 μm-thick sections through the barrel cortex, and 4-12 sections per brain (33-36 sections/genotype, 138 sections in total) were imaged with a 20X, 0.75 NA objective on a confocal microscope, with optical sections taken at 2.5 μm Z-steps. In each section, the imaged ROI was selected based on the extent of FB labeling (see below), and typically extended 700-1000 μm along the pial surface, spanning pia to white matter. The same ROIs (imaged with different laser lines) were used for quantifying fluorescent protein expression, FB labeling and immunostaining, as described below.

The pattern of fluorescent protein expression is illustrated in Fig. 1A by a representative confocal projection from each genotype. As evident from these images, the 4 genotypes had very different laminar distributions of GFP-expressing cells, with the Calb2 and Calb1 cell bodies distributed in L5 and L2/3, the Pdyn subset mostly in L4 and L5 and cell bodies of Chrna2 cells restricted to a narrow band in lower L5/upper L6. Also evident is a superficial band of axonal arborizations, restricted to L1 in the Chrna2 subset but encompassing also L2/3 in the Calb2 and Calb1 subsets, in addition to dense arborizations surrounding the cell bodies in L5. In the Pdyn genotype, axonal arborizations were most prominent in L4, in sharp contrast to the other three genotypes whose axons appeared to avoid L4.

The dual-color fluorescence of the combinatorial reporter allowed us to quantify not only the laminar distribution of neurons belonging to each subset, but also their prevalence within the overall SOM population. To do so, we counted the number of GFP-expressing and tdTomato-expressing cells by layer, by visual inspection of all optical planes imaged in each section. To correct for differences in the total cortical volume analyzed in different animals (e.g., due to variations in the extent of FB labeling), counts in each brain were normalized to all SOM cells (both GFP-expressing and tdTomato-expressing) counted in that brain, and then averaged within each genotype (Fig. 1B, upper panels; Table 1). For clarity, the same data are also shown further normalized to all SOM cells within each layer (Fig. 1B, lower panels). When grand-averaged over all 4 genotypes (***Table 1***, *last row*), about 15% of all SOM cells were found in L2/3, 10% in L4, 50% in L5 and 25% in L6. Calb2 cells comprised 10% of all SOM cells; they were found mostly in L2/3, where they represented nearly 40% of all SOM cells, with a minor population in L5 and very small numbers in L4 and L6. The Chrna2 group comprised 13% of all SOM cells straddling the L5/6 boundary, comprising ∼20% of L5 and ∼10% of L6 SOM cells. Calb1 cells were 63% of all SOM interneurons, ranging from 40-80% of SOM cells in different layers, with the lowest percentage in L4. Given that this subset includes 80% of all SOM cells in L2/3, it must contain at least half of all Calb2 cells in that layer. Lastly, Pdyn cells were slightly less than half of all SOM cells, making up 75% of SOM cells in L4 and about half of SOM cells in L2/3 and L5, with a small number in L6. Our counts for the Calb1 and Calb2 subsets are in excellent agreement with previous counts from immunostained brains of these genotypes (Nigro et al., 2018); our counts for the Pdyn subset are similar to, but slightly higher than counts from a different Pdyn-Cre intersection labeled by single-molecule FISH (Wu et al., 2022); and our counts of the Chrna2 subset are virtually identical to counts from a single-nucleus transcriptomic dataset of visual cortex (Wu et al., 2022).

**Table 1.** Laminar distributions of GFP expressing cells in the 4 intersectional subsets. For each genotype, the ***upper row*** indicates the number of GFP+ cells in each layer, as percentage of all SOM cells (GFP+ or tdTomato+) counted in each brain, averaged over all brains of the genotype. SEM indicated in parenthesis. The ***lower row*** expresses the same counts as percentage of all SOM cells in each layer. These data are plotted in **Fig. 1B**. The ***last row*** of the table quantifies the distribution of all SOM cells by layer, averaged over all 17 brains.

|  |  | **L2/3** | **L4** | **L5** | **L6** | **All layers** |
| --- | --- | --- | --- | --- | --- | --- |
| Calb2 (N=4) | % of all SOM | 6.0 (0.8) | 0.8 (0.3) | 2.3 (0.2) | 0.6 (0.2) | 9.7 (0.8) |
|  | % in layer | 36.1 | 6.6 | 4.8 | 2.6 | 9.7 |
| Chrna2 (N=6) | % of all SOM | 0.0 (0.0) | 0.0 (0.0) | 11.2 (0.5) | 2.2 (0.2) | 13.4 (0.5) |
|  | % in layer | 0.2 | 0.0 | 21.7 | 8.8 | 13.4 |
| Calb1 (N=4) | % of all SOM | 12.2 (0.5) | 4.5 (0.3) | 33.3 (1.2) | 12.9 (0.9) | 63.0 (1.6) |
|  | % in layer | 83.2 | 38.3 | 63.9 | 60.3 | 63.0 |
| Pdyn (N=3) | % of all SOM | 7.0 (1.3) | 9.3 (0.1) | 26.6 (2.8) | 3.8 (0.7) | 46.7 (4.6) |
|  | % in layer | 51.8 | 75.7 | 54.7 | 14.9 | 46.7 |
| All SOM (N=17) | % of all SOM | 14.4 (0.4) | 11.4 (0.3) | 50.5 (0.7) | 23.8 (0.6) | 100.0 |

### Overlap of intersectional subsets with transgenic subtypes

We previously developed and characterized two transgenic mouse lines, X98 and X94, with GFP expression restricted to specific subsets of SOM interneurons. X98 cells are L1-targeting (Martinotti) SOM neurons which reside mostly in the infragranular layers and could therefore overlap with Chrna2 neurons. X94 cells are L4-targeting (non-Martinotti) SOM neurons which reside in layers 4-5 and could therefore overlap with Pdyn neurons. To clarify the relationships between these subsets, we bred the hybrid genotypes Chrna2-Cre;X98 and Pdyn-Cre;X94 and crossed them with a Cre-dependent tdTomato reporter (Fig. 2A). We imaged fixed brain sections as described above and counted the fraction of X98 and X94 cells which also expressed tdTomato (Fig. 2B and Table 2). Less than one quarter of all X98 cells expressed tdTomato, indicating that the X98 and Chrna2 subsets are largely non-overlapping populations. In contrast, two-thirds of X94 cells expressed tdTomato and were thereby contained within the Pdyn subset.

**Figure 2.**
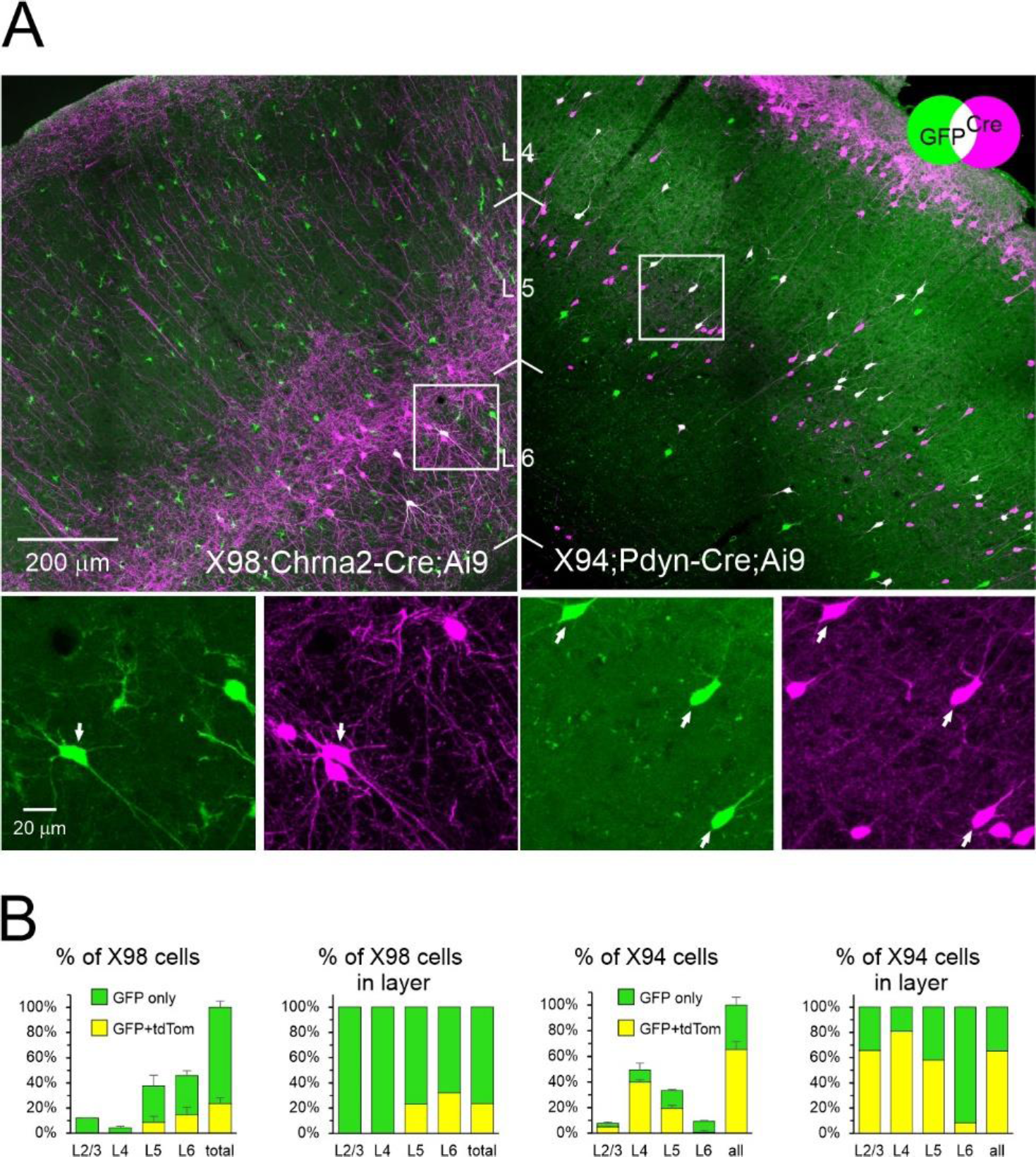
Overlap of intersectional and transgenic subsets. X98 mice were crossed with Chrna2-Cre and a tdTomato reporter, X94 mice were crossed with Pdyn-Cre and a tdTomato reporter. ***A,*** projections of confocal Z-stacks through representative 30 μm-thick sections of the indicated genotype, taken with a 20×, 0.75 NA objective at 2.5 μm Z-steps. Venn diagram illustrates the combinatorial color code. Color channels were adjusted individually. Boxed ROIs in ***upper panels*** are shown at higher magnification in ***lower panels***, separated into the two color channels. Arrows point to double-labeled cells. Note that a considerable population of L2 pyramidal neurons also expressed Pdyn-Cre, underscoring the need for the intersectional approach used here. ***B, left panel*** of each genotype shows fraction of GFP+ only and double-labeled cells (GFP+/TdTomato+) in each layer, out of all GFP+ cells counted in each brain, averaged by genotype. Error bars are SEM. The **right panel** of each genotype shows the same counts expressed as a fraction of all GFP+ cells in each layer. N=3, 2 for X98 and X94, respectively.

**Table 2.** Overlap between intersectional and transgenic SOM subsets. Counts are from the same crossed genotypes illustrated in Fig. 2. In each genotype, ***upper row*** is the distribution of GFP+ cells (i.e. X98 or X94) in each layer, as percentage of all GFP+ cells counted in each brain, averaged by genotype. SEM indicated in parenthesis. ***Middle rows***, double-labeled (DL, GFP+ and tdTomato+) cells in each layer, as percentage of all GFP+ cells counted in each brain, averaged by genotype. ***Lower rows***, the same numbers as percentage of all GFP+ cells in each layer. These data are also plotted in **Fig. 2B**.

|  |  | L2/3 | L4 | L5 | L6 | All layers |
| --- | --- | --- | --- | --- | --- | --- |
| X98;Chrna2-Cre (N=3) | % of all GFP+ | 12.1 (0.2) | 4.5 (1.1) | 37.7 (8.9) | 45.7 (8.7) | 100 |
|  | % DL of all GFP+ | 0.0 | 0.0 | 8.7 (4.8) | 14.7 (5.8) | 23.4 (4.9) |
|  | % DL in layer | 0.0 | 0.0 | 23.2 | 32.1 | 23.4 |
| X94;Pdyn-Cre (N=2) | % of all GFP+ | 7.8 (0.7) | 49.5 (3.6) | 33.4 (3.4) | 9.3 (0.9) | 100 |
|  | % DL of all GFP+ | 5.1 (1.0) | 40.0 (5.3) | 19.4 (1.0) | 0.8 (0.8) | 65.4 (6.2) |
|  | % DL in layer | 65.6 | 80.9 | 58.2 | 8.2 | 65.4 |

### Classifying SOM subtypes by their L1 projection

SOM interneurons with radially ascending axons projecting to L1 are historically referred to as Martinotti cells (Marin-Padilla, 1990; DeFelipe, 2002). Not all SOM interneurons, however, are L1-projecting; several important groups do not terminate in L1 and are collectively referred to as non-Martinotti (Tremblay et al., 2016). There is no quantitative estimate to-date on the proportion of anatomically verified Martinotti vs non-Martinotti SOM cells in the mouse cortex. To distinguish between Martinotti and non-Martinotti cells, we retrogradely labeled SOM neurons by placing a Fast Blue (FB)-infused filter paper on the pial surface, 24 hours prior to fixing the brains by transcardial perfusion (Cauller et al., 1998; Ramos-Moreno and Clasca, 2014). FB-labeled cells were then counted by visual inspection of 9-12 optical planes imaged through each tissue section. To validate this approach as a reliable method for labeling L1-projecting but not non-L1-projecting cells, we performed retrograde labeling in mice of the X98 and X94 lines, in which GFP-expressing SOM cells are L1-and L4-projecting, respectively (Ma et al., 2006). Overall, 84±5% (N=3) of all X98 cells, but only 14±3% (N=3) of all X94 cells (7% in L4) were FB-labeled (Fig. 3; Table 3), consistent with the distinct axonal projection targets of these transgenically defined SOM subsets.

**Figure 3.**
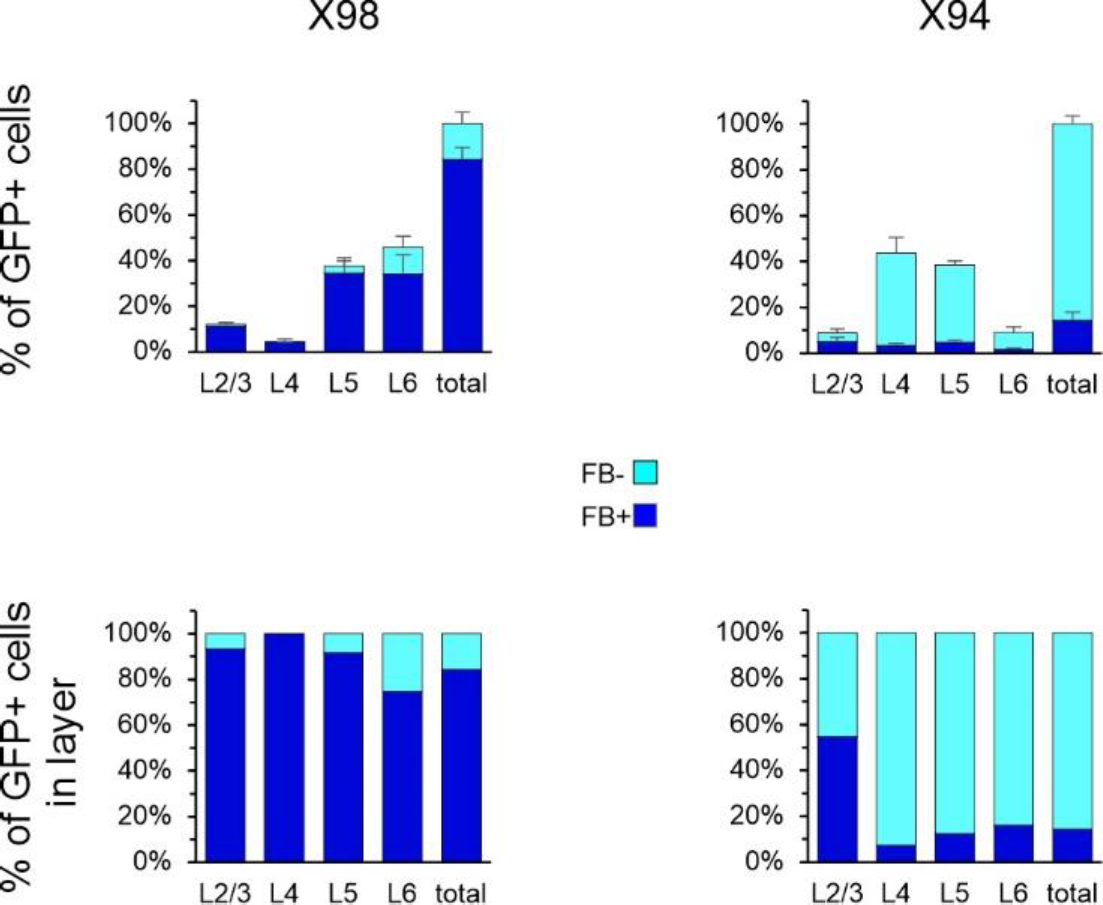
Retrograde FB labeling in the X98 and X94 subtypes. ***Top panels***, number of FB+ and FB-GFP+ cells, as a fraction of all GFP+ cells counted in each brain, averaged by genotype. ***Lower panels***, same numbers as a fraction of all GFP+ cells in each layer.

**Table 3.** Retrograde FB labeling in X98 and X94 mice. ***Upper rows*,** number of FB-labeled GFP+ cells per layer as percentage of all GFP+ cells counted in each brain, averaged by genotype. SEM indicated in parenthesis. ***Lower rows***, same numbers expressed as percentage of all GFP+ cells in each layer. These data are plotted in ***Fig. 3***.

|  |  | **L2/3** | **L4** | **L5** | **L6** | **All layers** |
| --- | --- | --- | --- | --- | --- | --- |
| X98 (N=3) | % FB+ of all GFP+ | 11.3 (0.9) | 4.5 (1.1) | 34.5 (6.8) | 34.1 (8.4) | 84.4 (5.1) |
|  | % FB+ in layer | 93.4 | 100.0 | 91.6 | 74.6 | 84.4 |
| X94 (N=3) | % FB+ of all GFP+ | 4.8 (2.0) | 3.2 (1.2) | 4.8 (0.9) | 1.4 (0.8) | 14.3 (3.5) |
|  | % FB+ in layer | 54.7 | 7.4 | 12.4 | 16.1 | 14.3 |

A confocal projection through a representative section from a retrogradely labeled Sst-Flp;Pdyn-Cre;RC::FLTG brain is illustrated in Fig. 4A, with the full cortical depth shown in the left panels and 4 selected ROIs (from L2/3-L6) magnified in the right panels. As seen in these images, the majority of FB-labeled cells in L2/3 and in L5 exhibited pyramidal morphology, and were most likely pyramidal cells labeled via their axonal terminations or dendritic tufts in L1. In contrast, L4 and L6 were mostly devoid of label, as excitatory neurons in these layers rarely extend dendrites or axons to L1 (Thomson, 2010; Oberlaender et al., 2012; Wang et al., 2018). Notably, a thin layer of cells abutting the subcortical white matter, in L6B (also referred to as L7, (Reep, 2000)), were found to be brightly labeled, as previously observed after pial dye deposits (Mitchell and Cauller, 2001; Ramos-Moreno and Clasca, 2014). This robust label in the deepest cortical layer indicated that the 24 hr survival time in our experiments was sufficient to retrogradely label any cortical neuron with an axonal projection in L1. In each section we characterized each GFP-expressing or tdTomato-expressing cell as either FB+ or FB-. The fraction of FB+ cells in each genetic subset is quantified by layer in Fig. 4B, both as a fraction of all cells of this subset (upper panels) and normalized by layer (lower panels) (Table 4). The majority of Calb2, Chrna2 and Calb1 cells were FB+ (70%, 90% and 64%, respectively), while only about half of the Pdyn subset were. In all subsets, nearly all FB+ cells were found in L2/3 and/or L5. Out of all SOM cells in all layers, 54±1% (N=17) were retrogradely labeled and therefore were bona-fide Martinotti cells. This fraction varied by layer, from 80% in L2/3 to ∼60% in L5, 40% in L6 and <20% in L4.

**Figure 4.**
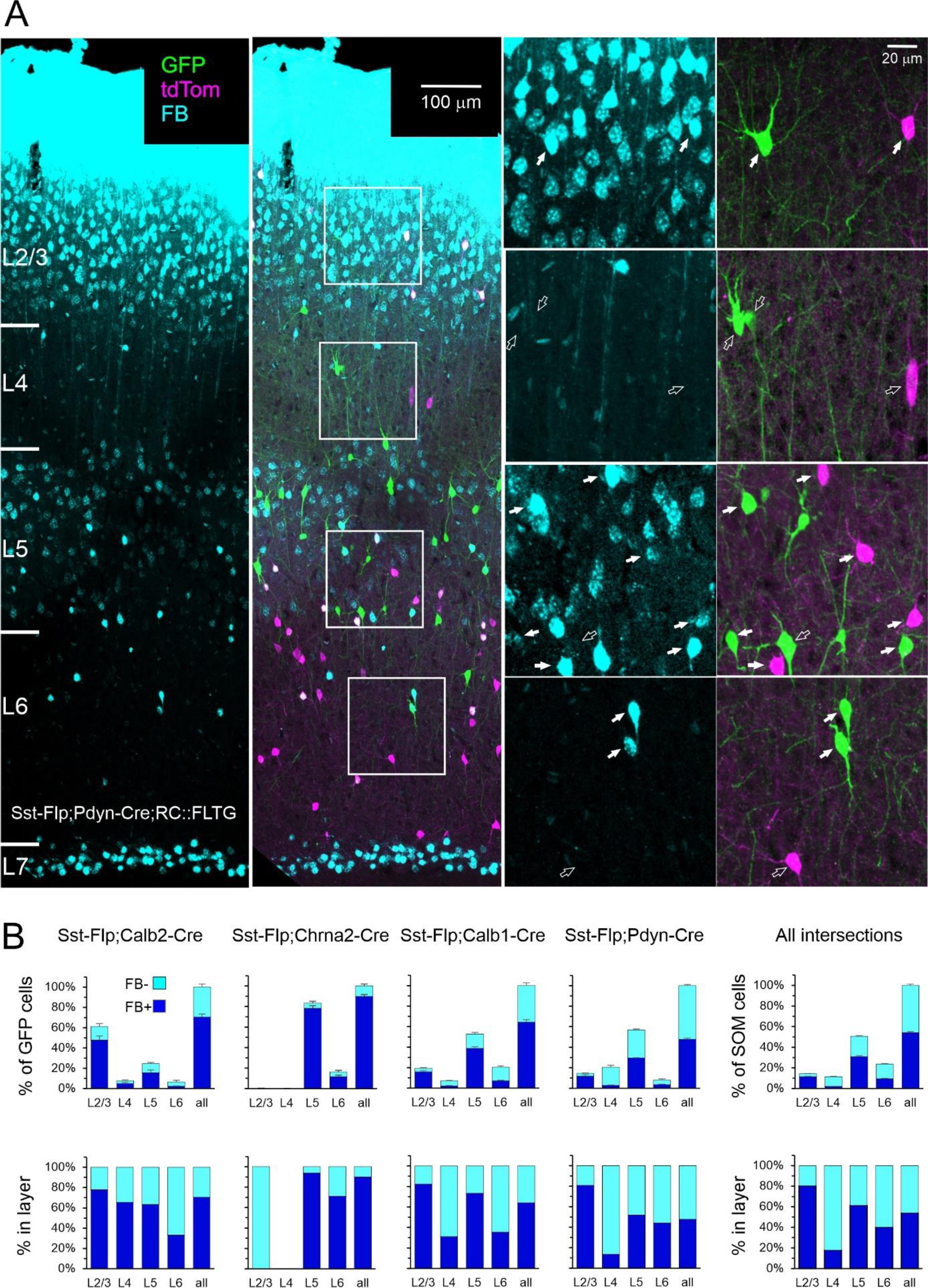
Retrograde FB labeling. ***A,*** representative section from a retrogradely labeled Pdyn mouse. Image is a projection of a Z stack taken with a 20×, 0.75 NA objective; color channels were adjusted individually. ***Left panel*** shows only the FB color channel, ***right panel*** also shows GFP and tdTomato channels. The 4 boxed ROIs are shown enlarged to the right, separated into the FB channel ***(left panels***) and GFP-tdTomato channels ***(right panels***). *Filled arrows* point to all FB+ SOM cells in each ROI, *hollow arrows* point to all FB-SOM cells. ***B, top panels***, number of FB+ and FB-GFP+ cells as percentage of all GFP+ cells counted in each brain, averaged by genotype. ***Lower panels***, same numbers as percentage of all GFP+ cells in each layer. Error bars are SEM. Plots at the far right show fraction of FB+ cells out of all SOM cells in all genotypes. Number of animals as in Fig. 1.

**Table 4.** Percentage of retrogradely labeled cells in the 4 intersectional subsets. In each subset, the ***upper row*** indicates number of FB-labeled GFP+ cells per layer as percentage of all GFP+ cells counted in each brain, averaged by genotype. SEM indicated in parenthesis. ***Lower row*** presents the same numbers expressed as percentage of all GFP+ cells in each layer. Empty cells indicate that no GFP+ neurons were found. The ***last 2 rows*** of the table quantify the distribution of all retrogradely labeled SOM cells (both GFP+ and tdTomato+), averages over all genotypes. These data are also plotted in **Fig. 4B**.

|  |  | **L2/3** | **L4** | **L5** | **L6** | **All layers** |
| --- | --- | --- | --- | --- | --- | --- |
| Calb2 (N=4) | % of all GFP+ | 47.7 (4.2) | 5.0 (1.1) | 15.5 (2.9) | 2.2 (0.5) | 70.5 (3.1) |
|  | % in layer | 78.0 | 65.5 | 63.4 | 33.2 | 70.5 |
| Chrna2 (N=6) | % of all GFP+ | 0.0 (0.0) |  | 78.3 (2.6) | 11.6 (1.5) | 89.9 (1.8) |
|  | % in layer | 0.0 |  | 93.8 | 71.1 | 89.9 |
| Calb1 (N=4) | % of all GFP+ | 16.0 (1.2) | 2.2 (0.3) | 38.7 (1.7) | 7.3 (0.4) | 64.2 (2.7) |
|  | % in layer | 82.2 | 31.0 | 73.3 | 35.4 | 64.2 |
| Pdyn (N=3) | % of all GFP+ | 11.9 (1.5) | 2.8 (0.4) | 29.5 (0.3) | 3.6 (0.3) | 47.7 (1.2) |
|  | % in layer | 80.8 | 13.6 | 51.9 | 44.2 | 47.7 |
| All SOM (N=17) | % of all SOM | 11.5 (0.3) | 2.0 (0.2) | 30.9 (0.9) | 9.5 (0.3) | 54.0 (1.2) |
|  | % in layer | 80.3 | 17.9 | 61.3 | 40.0 | 54.0 |

### Expression of protein markers in SOM subtypes

Cortical interneurons can be differentiated by their pattern of expression of a variety of calcium-binding proteins and neuropeptides (Demeulemeester et al., 1988; Baimbridge et al., 1992; Kubota et al., 1994). We tested the 4 intersectional subsets for immunostaining against three proteins known to be expressed by SOM interneurons in the mouse: calretinin (CR, product of the Calb2 gene), neuropeptide Y (NPY) and calbindin (CB, product of the Calb1 gene) (Xu et al., 2010). Each antibody was tested on (typically) 4 sections/brain from 3 brains/genotype; only one antibody was tested on each section. Three representative sections stained with the 3 antibodies, respectively, are shown in Fig. 5A, with one L5 ROI in each section shown enlarged to the right of the low-power image.

**Figure 5.**
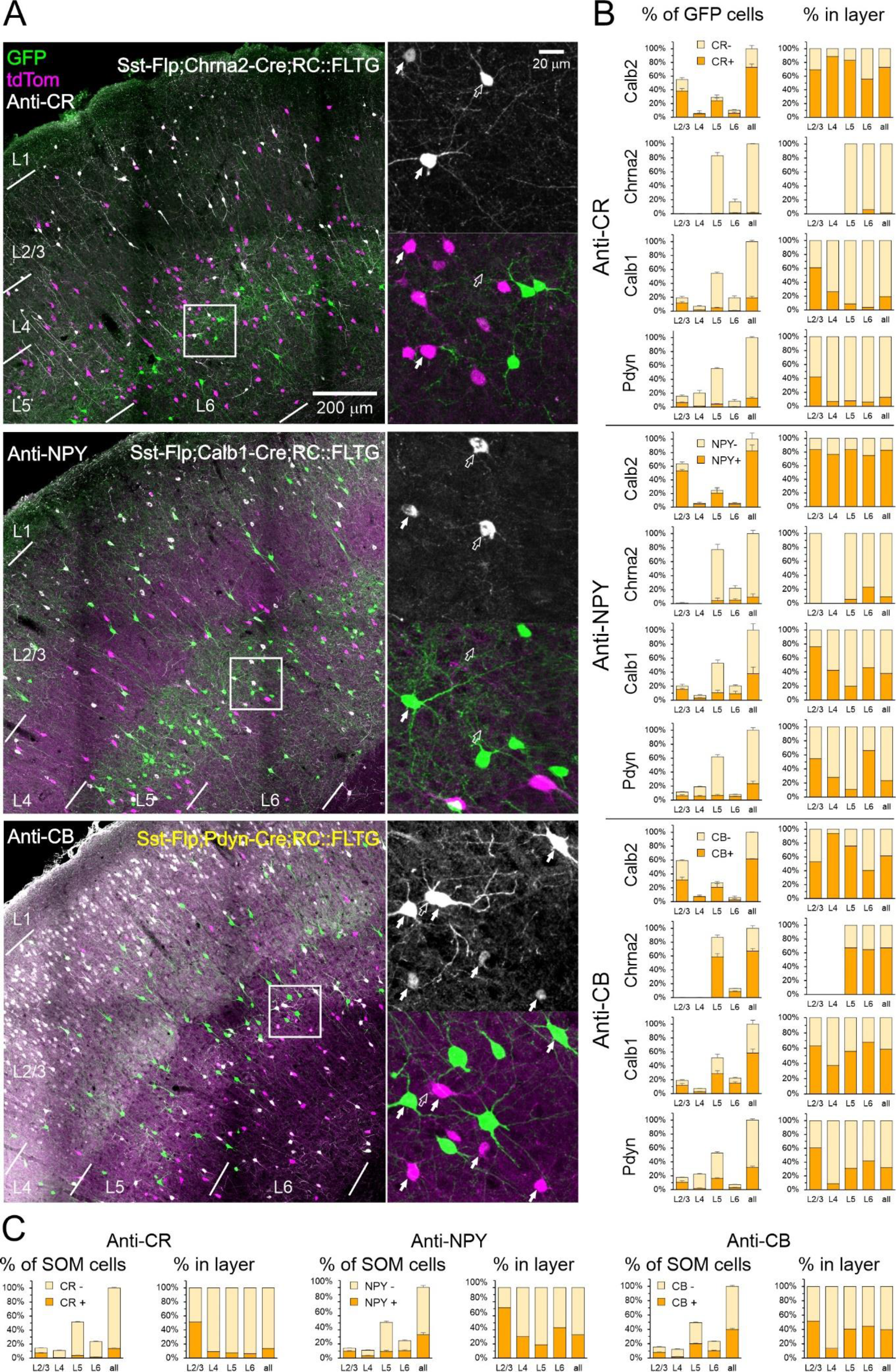
Protein marker expression in the 4 intersectional subsets. ***A, upper panel***, anti-CR immunostaining on a Chrna2 section. ***Middle panel*,** anti-NPY immunostaining on a Calb1 section. ***Lower panel***, anti-CB immunostaining on a Pdyn section. All images are projections of Z stacks taken with a 20×, 0.75 NA objective. Color channels were adjusted individually. In each section, one ROI is shown enlarged in the right panels, separated by color channels. Arrows point to all immunopositive cells within the ROI; filled arrows indicate SOM cells (GFP+ or tdTomato+), hollow arrows indicate non-SOM cells. ***B, left panels,*** percentages of CR, NPY, and CB immunostained cells out of all GFP+ cells counted in each brain, averaged by genotype; error bars are SEM. ***Right panels,*** same numbers expressed as percentages of all GFP+ cells counted in each layer. N=3 mice for each antibody. ***C,*** same counts expressed as percentages of all SOM cells (GFP+ and tdTomato+) in each brain, averaged over all 4 subsets, N=12.

As for FB, we characterized each GFP or tdTomato-expressing cell as positive or negative for the antibody tested on that section, by examining all optical planes taken through the section (see high-power panels in Fig. 5A, right column). For each intersectional subset, we quantified the fraction of all GFP-expressing neurons labeled by each antibody in each layer, averaged over the 3 brains. These counts are plotted in Fig. 5B, left panels. The same data are also plotted in the right panels normalized to the average count in each layer, for clarity. We also quantified the fraction of all SOM cells (both GFP and tdTomato-expressing) immunostained by each antibody, averaging over all 12 brains. These data are plotted in Fig. 4C in the same manner as Fig. 4B, with the right plot in each pair of plots showing the averaged counts normalized by layer. These data are also provided in Table 5.

**Table 5.** Marker protein immunoreactivity in the 4 intersectional subsets. For each antibody, ***upper row*** indicates count of immunostained cells in each layer, expressed as percentage of all GFP+ cells in the same sections; SEM indicated in parenthesis. *L**ower row*** expresses the same counts as percentage of all GFP+ cells in the same layer. *Empty cells* indicate that no GFP+ neurons were found. N=3 mice for each antibody in each genotype; most brains were used for 2-3 antibodies each. These data are plotted in ***Fig. 5B***. The ***last block*** in the table indicates counts of immunostained cells expressed as percentage of all SOM cells (GFP+ and tdTomato+) counted in each brain, averaged over all genotypes, N=12 mice per antibody. These data are plotted in ***Fig. 5C***.

|  |  | **L2/3** | **L4** | **L5** | **L6** | **All layers** |
| --- | --- | --- | --- | --- | --- | --- |
| **Calb2 (N=3)** |  |  |  |  |  |  |
| CR+ | % of all GFP+ | 38.2 (3.7) | 4.8 (4.8) | 24.3 (3.7) | 5.7 (1.8) | 73.1 (4.6) |
|  | % in layer | 69.3 | 88.9 | 83.2 | 55.9 | 73.1 |
| NPY+ | % of all GFP+ | 53.1 (2.0) | 4.4 (1.5) | 20.6 (6.5) | 4.4 (1.5) | 82.5 (8.6) |
|  | % in layer | 83.4 | 76.5 | 83.5 | 75.0 | 82.5 |
| CB+ | % of all GFP+ | 31.3 (4.0) | 7.1 (2.6) | 20.7 (5.3) | 2.4 (1.8) | 61.5 (0.5) |
|  | % in layer | 53.0 | 93.9 | 75.8 | 40.4 | 61.5 |
| **Chrna2 (N=3)** |  |  |  |  |  |  |
| CR+ | % of all GFP+ |  |  | 0.4 (0.4) | 1.0 (0.6) | 1.4 (0.7) |
|  | % in layer |  |  | 0.5 | 6.0 | 1.4 |
| NPY+ | % of all GFP+ | 0.0 (0.0) |  | 4.3 (3.7) | 5.1 (2.0) | 9.4 (4.7) |
|  | % in layer | 0.0 |  | 5.6 | 23.3 | 9.4 |
| CB+ | % of all GFP+ |  |  | 58.6 (4.7) | 8.5 (1.2) | 67.1 (3.7) |
|  | % in layer |  |  | 67.4 | 64.9 | 67.1 |
| **Calb1 (N=3)** |  |  |  |  |  |  |
| CR+ | % of all GFP+ | 11.7 (1.7) | 1.9 (0.3) | 4.8 (0.6) | 0.8 (0.3) | 19.2 (2.3) |
|  | % in layer | 61.0 | 26.4 | 8.9 | 4.0 | 19.2 |
| NPY+ | % of all GFP+ | 15.3 (1.6) | 2.9 (0.7) | 10.4 (3.8) | 9.4 (3.3) | 38.0 (9.2) |
|  | % in layer | 76.2 | 42.5 | 19.7 | 46.3 | 38.0 |
| CB+ | % of all GFP+ | 11.9 (2.8) | 2.7 (0.6) | 28.7 (4.2) | 15.2 (1.8) | 58.6 (5.6) |
|  | % in layer | 62.9 | 37.4 | 55.9 | 67.9 | 58.6 |
| **Pdyn (N=3)** |  |  |  |  |  |  |
| CR+ | % of all GFP+ | 6.6 (0.9) | 1.4 (0.3) | 4.4 (0.5) | 0.5 (0.3) | 13.0 (1.5) |
|  | % in layer | 42.3 | 7.0 | 8.0 | 6.1 | 13.0 |
| NPY+ | % of all GFP+ | 6.3 (1.3) | 5.3 (1.3) | 6.7 (1.4) | 5.2 (0.5) | 23.5 (3.6) |
|  | % in layer | 54.6 | 28.1 | 10.9 | 66.3 | 23.5 |
| CB+ | % of all GFP+ | 10.6 (2.3) | 2.0 (0.4) | 16.4 (0.7) | 3.1 (0.4) | 32.0 (2.0) |
|  | % in layer | 60.6 | 8.7 | 31.1 | 41.6 | 32.0 |
| **All SOM** |  |  |  |  |  |  |
| CR+ (N=12) | % of all SOM | 7.3 (0.6) | 1.0 (0.2) | 3.8 (0.3) | 1.5 (0.1) | 13.6 (0.9) |
|  | % in layer | 51.5 | 9.0 | 7.4 | 6.4 | 13.6 |
| NPY+ (N=12) | % of all SOM | 9.7 (1.0) | 3.4 (0.6) | 9.4 (1.4) | 10.7 (0.9) | 33.3 (3.0) |
|  | % in layer | 71.4 | 30.6 | 18.7 | 43.3 | 33.3 |
| CB+ (N=12) | % of all SOM | 7.9 (0.6) | 1.7 (0.2) | 20.0 (0.9) | 10.3 (0.6) | 39.9 (1.4) |
|  | % in layer | 51.6 | 13.8 | 40.5 | 44.5 | 39.9 |

In the Calb2 subset, about 75% of neurons were CR+, with some minor variations between layers, as reported in a previous study of this intersection (Nigro et al., 2018). That not all Calb2 cells expressed CR, even though Cre and CR proteins were presumably translated from a single bicistronic transcript, could mean that some neurons expressed CR and Cre during prenatal or early postnatal development, underwent Cre-mediated recombination but no longer expressed the marker protein in the adult animals tested here. The majority (∼60%) of Calb2 cells were also immunopositive for CB, a fraction very similar to that of triple-labeled CB+, CR+, SOM+ cells out of all CR+, SOM+ cells (Nigro et al., 2018). An even larger fraction of Calb2 cells (∼80%) were NPY+, suggesting that most Calb2 cells expressed both CR and NPY. High co-expression of CR and NPY in upper layers SOM cells was previously noted in the cingulate cortex (Riedemann et al., 2016).

In contrast to Calb2 cells, 2/3 of Chrna2 SOM cells were immunopositive to CB but only 10% to NPY and nearly none to CR, indicating that the Calb2 and Chrna2 subsets were distinct populations with little or no overlap. In the Calb1 subset, 20, 40 and 60% of cells were immunopositive for CR, NPY and CB, respectively. Lastly, Pdyn SOM cells were immunopositive to the 3 markers at about half the rates of Calb1 cells (13%, 23% and 32% for CR, NPY and CB, respectively). Especially low levels (<10%) of CR and CB immunoreactivity were found in L4 Pdyn cells, consistent with the absence of these markers from L4 X94 neurons (Ma et al., 2006; Naka et al., 2019).

In all SOM cells (Fig. 5C), the highest incidence of the 3 markers was found in L2/3, where at least 50% of SOM cells were immunopositive for each marker (tested separately). Fractions were lower in the other layers, especially for CR, detected in less than 10% of SOM cells in each of the other layers. A very similar laminar distribution of CR+ SOM cells in mouse S1 was reported previously by Xu et al., 2010. Notably, however, our counts for NPY in SOM cells (70% immunopositive in L2/3, 33% overall) were several fold higher than reported by Xu et al. (∼10% and 7%, respectively). This discrepancy could reflect the low sensitivity of the anti-NPY antibody used in the Xu et al. study. Indeed, in a previous study by the same authors (Xu et al., 2006) using the same antibody, no NPY expression was detected in GIN cells, a subset of GFP-expressing SOM interneurons in L2/3 and L5, whereas two previous studies (in S1 and in cingulate cortex), using two NPY antibodies different from the ones used by Xu et al. and in the current study, found that about 30% of L2/3 GIN cells were NPY+ (Ma et al., 2006; Riedemann et al., 2016). That NPY expression was strongly genotype dependent – e.g. >80% of all Calb2 cells, but <10% of Chrna2 cells were immunopositive for NPY – also indicated that our NPY antibody was both sensitive and specific. Lastly, a recent study using the same NPY antibody used here found that 50% of all SOM cells in adult mouse S1 cortex were NPY+ (Asgarian et al., 2022). We conclude that the NPY antibody we used provided an accurate estimate of NPY expression in SOM interneuron subsets.

### Multidimensional analysis of SOM subtypes

Since retrograde labeling and immunocytochemical staining were done on the same brains, we could relate the genetic label of a neuron (as a member of one of the 4 intersectional subsets) both to its axonal projection and to its protein marker expression. To convey these multi-dimensional data graphically, we present them as a grid of “sunburst charts” (Figure 6), with each row representing one subset and each column representing one of the 3 antibodies. Charts consist of 4 concentric rings, each showing the fractional value of one dimension of the data as a colored sector, normalized to all SOM cells and averaged over 3 brains. From the center out, the rings correspond to cortical layer, fluorescent protein expression, FB label and immunolabel. The inner ring is divided into 4 sectors, and each sector is further split two-way with each consecutive ring, so the outer ring consists of 32 sectors representing unique combinations of laminar position, fluorescent protein expression, FB label and antibody staining status. In addition to providing a succinct visual summary of the data already presented separately in Figs. 1B, 4B and 5B, ***Fig. 6*** demonstrates how different characteristics (such as retrograde labeling and immunostaining) converge on the same neurons, and illustrates counts of both GFP+ and tdTomato+ cells in each genotype. The full dataset is provided as extended data in ***Fig. 6-1***. An interactive version of these plots can be accessed at https://sites.google.com/view/somatostatinsubtypes/home.

**Figure 6.**
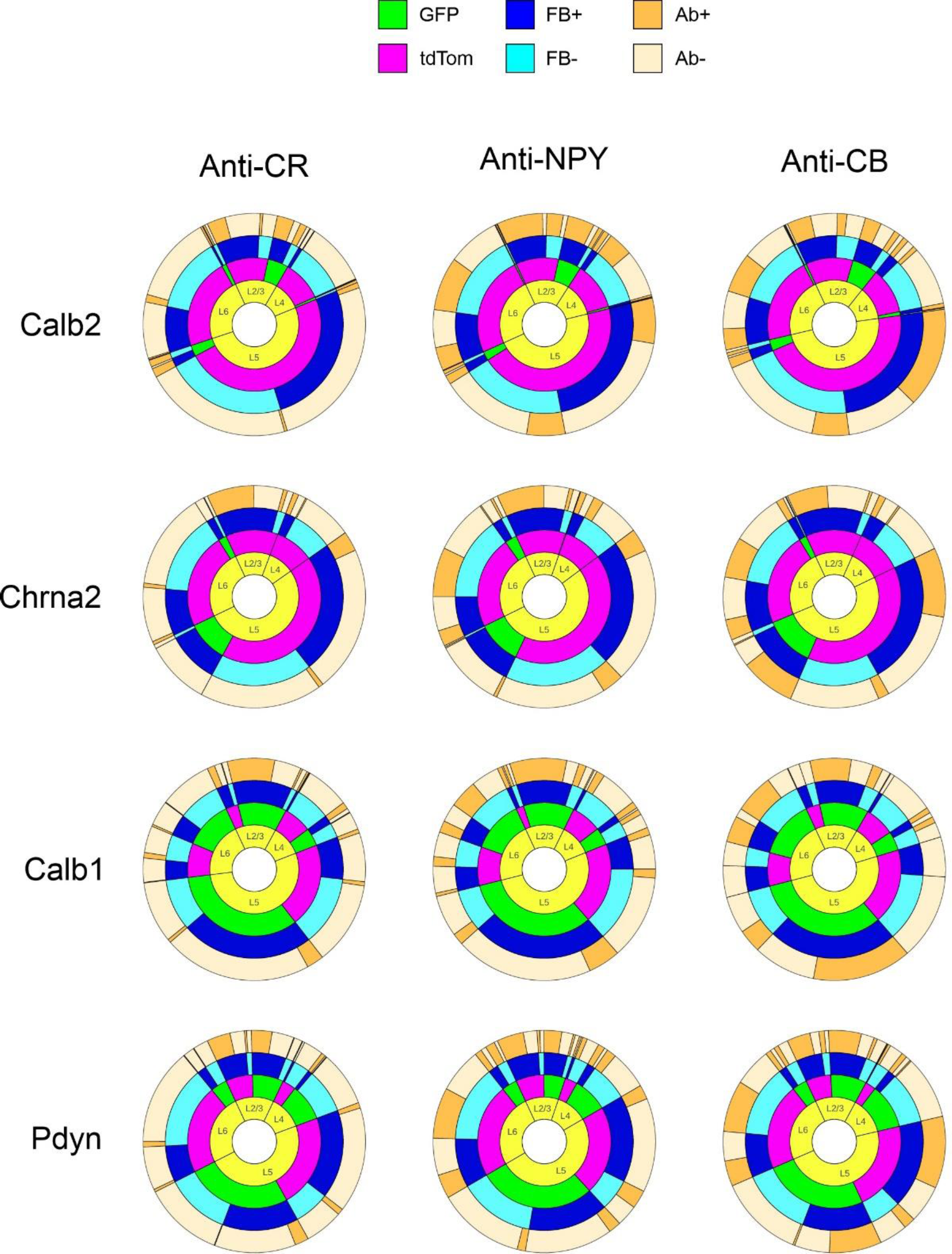
Sunburst charts representing fractional counts of the combined label (fluorescent protein, FB and antibody) of SOM cells in each genotypes for each antibody. N=3 brains per chart. See text for detailed explanation of sunburst charts. The data used to generate these charts can be found in ***Extended Data, Fig. 6-1***. An interactive version can be accessed at https://sites.google.com/view/somatostatinsubtypes/home

We can use ***Fig. 6*** to summarize the characteristics of each of the 4 intersectional subsets. Calb2 cells (green sectors in top row) were located mostly in L2/3, with a smaller population in L5. In both layers most Calb2 cells were L1-projecting (dark blue sectors) and expressed CR, NPY, and CB (dark tan sectors). Chrna2 cells were located exclusively in L5/6, and nearly all were L1-projecting. They expressed CB but, unlike Calb2 cells, almost never NPY or CR, and therefore these two subsets are non-overlapping SOM populations. Calb1 cells comprised the majority of SOM cells in L2/3, L5 and L6; nearly all in L2/3, and the majority in L5, were L1-projecting. Most L2/3 cells in this subset were positive for all 3 markers, while in L5 about half expressed CB but only a small minority expressed CR or NPY. Pdyn cells were located in all layers, comprising at least half of all SOM cells in L2/3, L4 and L5. Most of L2/3 Pdyn cells were L1-projecting, as indeed were most other SOM cells in this layer; however only about half of L5 Pdyn cells, and nearly none in L4, were L1-projecting, consistent with the overlap of this subset with L4-targeting (non-Martinotti) X94 cells in these layers. The majority of Pdyn SOM cells in L2/3 were positive for each of the 3 protein markers, but nearly none of those in L4 or in the FB-negative sector in L5 were immunopositive, consistent with the known absence of any of the marker proteins in X94 cells (Ma et al., 2006; Naka et al., 2019). We conclude from our multidimensional data that the Calb2 and Chrna2 subsets are relatively small, homogeneous and distinct groups, while the Calb1 and Pdyn subsets are larger (each comprising close to, or about half, of all SOM interneurons) and non-homogeneous. Both contain L1-projecting cells in L2/3, most of which express all 3 markers, and also contain L1-projecting cells in L5, most of which express CB. The Calb1 subset likely encompasses all or most of the Calb2 group, and the Pdyn group is likely a superset of the non-Martinotti X94 subtype.

### Electrophysiological classification of SOM subtypes

SOM interneurons have been shown to exhibit a diversity of electrophysiological properties, allowing (in some studies) their parcellation into subtypes which also exhibit distinct morphological and neurochemical phenotypes, or distinct gene expression patterns (Halabisky et al., 2006; Ma et al., 2006; McGarry et al., 2010; Riedemann et al., 2018; Gouwens et al., 2020). Given the clear separation between the X94, Chrna2 and Calb2 subsets based on neurochemical markers, it was therefore of interest to examine their electrophysiological phenotypes, and to test how well they can be classified based on these properties. Also, given the high prevalence of non-Martinotti L4 and L5 cells in the Pdyn subset, it was important to compare electrophysiological characteristics of Pdyn neurons to those of the previously characterized X94 subtype (Ma et al., 2006). We therefore recorded ex-vivo from GFP-expressing Calb2 (N=16 cells in 9 animals), Chrna2 (N=38 cells in 21 animals), Pdyn (N=23 cells from 6 animals, including one Pdyn-Cre;X94;Ai9 mouse) and X94 neurons (N=17 cells from 8 animals), focusing on L5 where all four subsets overlap. Animals were (typically) 1-2 months old, of both sexes. We did not include Calb1 neurons in these recordings because our histological analysis indicated that this genotype captures unselectively nearly all L1-targeting subtypes.

X94, Chrna2 and Calb2 SOM cells had distinct firing patterns (***Fig. 7, upper panels***). While all subsets exhibited pronounced spike-rate adaptation during a 600 ms suprathreshold current step, each had unique features which distinguished it from the other two. X94 cells, previously characterized as “quasi fast-spiking” (Ma et al., 2006), had considerably faster (narrower) spikes and higher steady-state firing frequency compared to the other two subsets. Chrna2 neurons fired a characteristic low-threshold burst at the onset of the current step, and as previously reported (Hilscher et al., 2017) also fired a burst upon recovery from hyperpolarization; 87% of all Chrna2 cells fired a burst, but only 13% of Calb2 cells, and no X94 did. Chrna2 cells also had the highest values of input resistance (note low-amplitude current steps). Calb2 cells fired at considerably lower rates compared to the other two subsets, a difference which was especially pronounced at the beginning of the current step, before spike frequency adaptation took place. Unlike the relatively homogeneous properties of these 3 subsets, Pdyn neurons exhibited a dichotomy: some had X94-like firing patterns and spike waveforms, and the rest resembled Chrna2 neurons (***Fig. 7, lower panels***), suggesting that this group included neurons from both the X94 and the Chrna2 subsets. While our histological analysis suggested that the Pdyn group is likely to contain also Calb2 cells (based on NPY and CR immunostaining), we did not encounter Pdyn cells with Calb2-like electrophysiological phenotype, possibly because Calb2 cells are a small minority (5%) of L5 SOM cells (***Fig. 1B***).

**Fig. 7:**
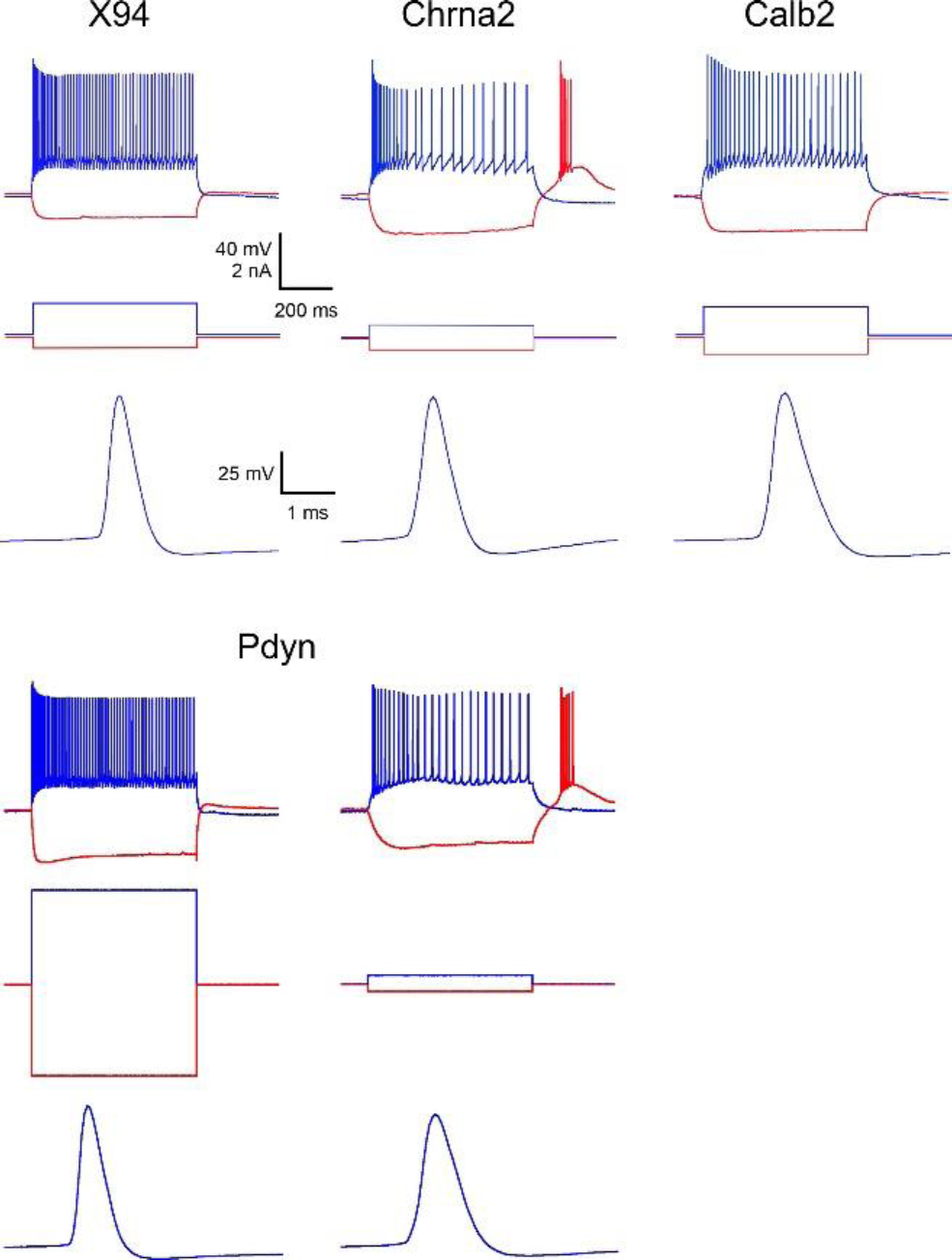
Characteristic firing patterns of the 4 subtypes. For each genotype, the ***upper panel*** illustrates response to a suprathreshold current step (blue trace) superimposed on a hyperpolarizing response to a negative current step (red trace); ***middle panel*** shows the applied current steps; and ***lower panel*** shows waveform of a single action potential evoked at rheobase, at an expanded time scale. Note the narrow spike and high steady-state firing frequency of the X94 neuron; the rebound burst and high input resistance of the Chrna2 cell; and the low firing frequency of the Calb2 cell. The Pdyn subset displayed a mix of the X94 and Chrna2 patterns.

To arrive at an objective classification of L5 SOM cells by their electrophysiological properties, we quantified 10 intrinsic electrophysiological properties for each cell, in addition to rebound spiking (see Methods). Other than spike height and sag, all parameters were significantly different between groups (permutation test on the F statistic), with 4 of them – input resistance (R_in_), spike width (SWHH), maximal firing frequency (F_max_) and steady-state firing frequency (F_SS_) – different at the p<0.0001 level. A plot of F_max_ vs R_in_ (***Fig. 8, left panel***) revealed a clear separation with only minor overlap between the 3 subsets. We then applied to this dataset two different dimensionality reduction methods – principal component analysis (PCA) and discriminant function analysis (DFA) (Manly, 2005; Ma et al., 2006; Druckmann et al., 2013). PCA is agnostic to the categorical identity (genotype) of each cell, and resulted in only minor improvement in the segregation of datapoints into 3 clusters (***Fig. 8, middle panel***). DFA is designed to maximize the separation between pre-categorized groups, and resulted in an almost complete segregation of the three clusters (***Fig. 8, right panel***). The two Calb2 cells intermixed among the Chrna2 datapoints in all 3 plots (designated by filled circles) were the only two Calb2 cells to display rebound bursting. Including the Pdyn subset in this analysis resulted in Pdyn datapoints scattered within both Chrna2 and X94 point clouds (not shown). The full dataset is available as ***Extended Data, Fig. 8-1***.

**Fig. 8:**
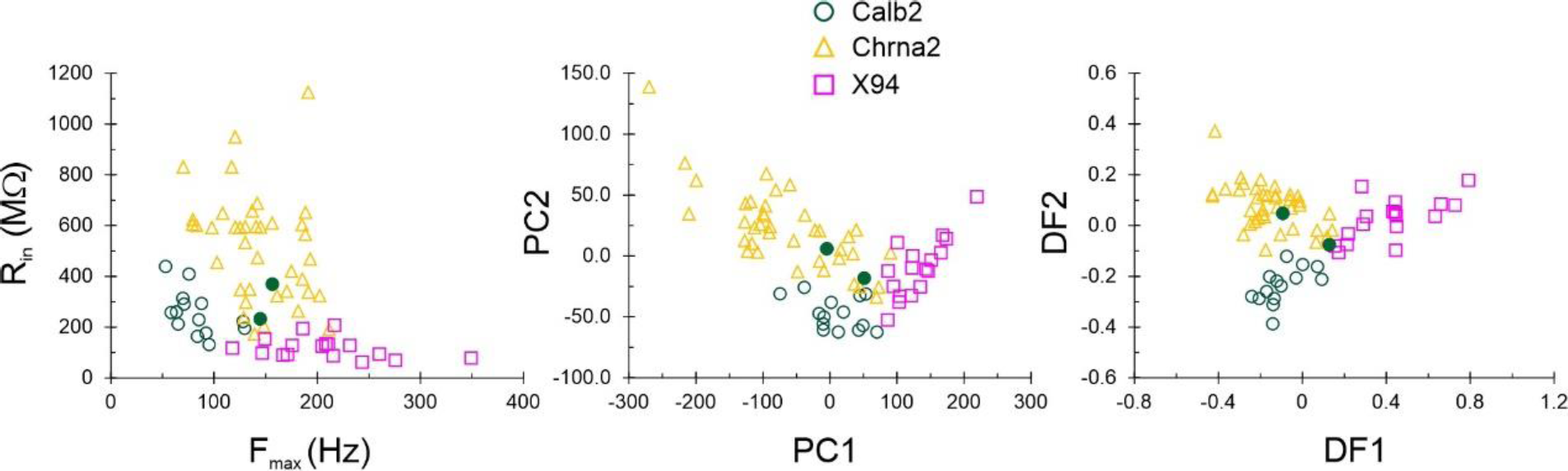
Multivariate analysis of electrophysiological properties in 3 genetically defined SOM subtypes. ***Left***, plot of maximal firing frequency vs input resistance separates the 3 subtypes. ***Middle*,** Principal component analysis leaves some overlap between the Chrna2 and Calb2 point clouds. ***Right***, Discriminant function analysis results in near-perfect segregation of the 3 subtypes. The two datapoints designated by filled circles are the only two Calb2 cells which exhibited rebound spiking. The full dataset is provided in ***Extended Data, Fig. 8-1***.

Previous machine-learning approaches found classification of cell types by electrophysiological properties alone challenging. For example, using a set of 44 electrophysiologically derived parameters, a random forest classifier achieved only 60% accuracy in assigning neurons to their correct transcriptomic type (Gouwens et al., 2020), and a K-nearest-neighbor classifier was unable to correctly separate Chrna2 from Calb2 neurons (Wu et al., 2022). To test how well electrophysiological properties can predict cell subtype, we constructed a simple decision tree with 5 nodes (decision points), using as input the 4 parameters different at the p<0.0001 level (***Fig. 9***). The first 3 nodes classified correctly all 17 X94 cells, but misclassified one Chrna2 cell with exceptionally narrow spikes as X94. The last two nodes classified correctly all but one of the remaining 37 Chrna2 cells (which was misclassified as Calb2), and all but two of the 16 Calb2 cells (misclassified as Chrna2). Interestingly, even though rebound bursting was not used as a decision parameter, the misclassified Chrna2 cell was among the only 5 of its group which did not burst, and the two misclassified Calb2 cells were again the only ones of their group to exhibit rebound bursts (filled circles in ***Fig. 8***). We conclude that the genetically defined SOM subtypes we studied here can be reliably and accurately classified based on a small number of basic electrophysiological properties.

**Fig. 9.**
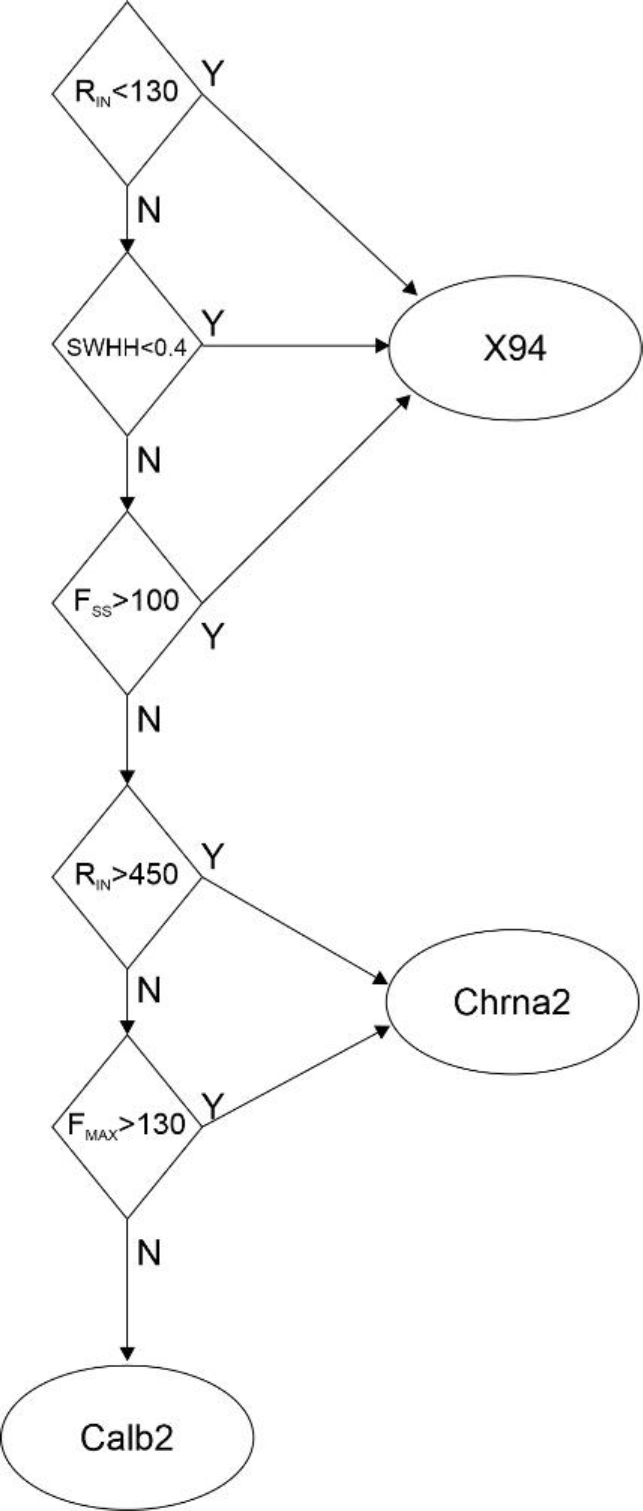
A decision tree for classifying L5 SOM subtypes by four basic electrophysiological properties. See Methods for parameter definitions. Units are Rin: [MΩ], SWHH: [ms], Fmax and Fss: [Hz].

## DISCUSSION

In an earlier study (Ma et al., 2006) we identified two SOM subtypes, based on GFP expression in the X94 and X98 transgenic lines. X98 cells were L1-targeting, had high input resistance and relatively slow spikes, fired low-threshold spike bursts, and expressed calbindin and (variably) NPY. X94 cells were L4-targeting, had lower input resistance and faster spikes, fired at high frequencies and did not express these protein markers. More recently, Hilscher et al. (Hilscher et al., 2017) described Chrna2 neurons in L5 as L1-targeting cells which fire low-threshold bursts; in this they resembled X98 cells, but no marker proteins (other than somatostatin) were tested. An in-vivo recording study (Munoz et al., 2017) identified two morphological types of L1-projecting SOM cells in L5: “T-shaped”, with a single main axon extending to L1 before branching, and “fanning-out”, with multiple ascending axon collaterals and with dense arborizations in both L2/3 and L1, in addition to L4-projecting non-Martinotti SOM cells. These three morphological types also exhibited distinct behaviorally linked activity patterns. A subsequent ex-vivo study (Nigro et al., 2018) observed the same 3 morphological types and found “fanning-out” cells preferentially among the Calb2 subset.

The studies above were conducted by three different laboratories using disparate methods, and left several question unanswered. Are the Chrna2, Calb2 and Calb1 subsets disjoint or overlapping? What protein markers are differentially expressed between them? What fraction of the L5 SOM population is captured by each of them? How are these 3 genotypes related to the transgenic X94 and X98 subsets? And finally, how do these different SOM subsets map onto the recent multimodal MET taxonomy, developed by yet another group of investigators? Here we set out to address these questions by examining all 5 previously studied genotypes and one novel genotype side-by-side. We found a clear separation between three SOM subsets - Chrna2, Calb2 and X94 – which constitute non-overlapping populations and differ from each other in both categorical and quantitative properties. As summarized graphically in ***Fig. 6***, the Chrna2 subset resides in L5b and Calb2 neurons are split 2:1 between L2/3 and L5/6. Both subsets are L1-projecting (Martinotti) neurons and express CB, but unlike Chrna2 cells, most Calb2 cells also express CR and NPY, indicating that these two subsets are fully disjoint. These characteristics also set them apart from X94 cells, which reside in L4/5, are L4-projecting and do not express any of these protein markers (Ma et al., 2006; Naka et al., 2019). Moreover, these three genetically-defined subsets have distinct firing patterns (***Fig. 7***) and segregate into non-overlapping clusters by their electrophysiological parameters (***Figs. 8,9***). Taken together, this multimodal correspondence of axonal target, neurochemical markers and electrophysiological properties (summarized in ***Table 6***) strongly supports consideration of these 3 genetically-defined subsets as bona-fide neuronal subtypes of the SOM subclass.

**Table 6.** Comparison of the main characteristics of the 3 L5 SOM subtypes. Approximate IQR (inter-quartile range, 25^th^-75^th^ percentiles) is listed for each of the 4 basic parameters used in Fig. 9 for classification. IQRs were largely non-overlapping between subsets, except for overlap of SWHH and of FSS between Chrna2 and Calb2.

|  | Laminar position | Axonal target | Protein markers | Electrophysiological characteristic | SWHH IQR (ms) | R<sub>in</sub> IQR (MΩ) | F<sub>max</sub> IQR (Hz) | F<sub>ss</sub> IQR (Hz) |
| --- | --- | --- | --- | --- | --- | --- | --- | --- |
| X94 | L4, L5 | L4 | ----- | Quasi fast-spiking | 0.35-0.45 | 90-140 | 170-230 | 90-120 |
| Chrna2 | L5/6 | L1 | CB | Low-threshold bursts | 0.45-0.6 | 340-600 | 120-180 | 35-60 |
| Calb2 | L2/3, L5 | L1 | CB,CR,<br>NPY | Low F<sub>max</sub> | 0.5-0.8 | 200-300 | 70-100 | 35-60 |

It is informative to estimate how many of all SOM cells, and specifically of L5 SOM cells, belong to these three largely disjoint subtypes. The Calb2 and Chrna2 groups together account for 27% of L5 SOM cells and 23% of all SOM cells (***Table 1***). To estimate the size of the X94 subset, we assume that it consists of all FB-negative Pdyn cells in L4 and L5, which comprise 26% of L5 SOM cells and 21% of all SOM cells (***Extended Data, Fig. 6-1***). This assumption may be an underestimate, as some X94 cells may be retrogradely labeled by FB (14% of X94 cells in all layers, ***Fig. 3***), but it may also be an overestimate, as some non-L1-targeting Pdyn cells in L5 may belong to other subtypes. Keeping these uncertainties in mind, we estimate that the three subtypes together account for about half of L5 SOM cells, and for >40% of all SOM interneurons.

How do these three SOM subtypes fit within the recent multimodal MET classification (Gouwens et al., 2020)? The L2/3 and L5 members of the Calb2 subtype appear to correspond to Sst-MET types 3 and 4, respectively, based on CR expression, L1-targeting and laminar position; some L2/3 Calb2 cells may also be included in Sst-MET 2. Chrna2 cells likely correspond to Sst-MET type 6, based on Chrna2 expression, L1-targeting and laminar position. Interestingly, the Sst-MET 5 type expresses lower levels of Chrna2, but shares with Sst-MET 6 high expression of CoupTFII/Nr2f2, and contains L5 neurons with a pronounced bursting phenotype (Gouwens et al., 2020). We hypothesize that X98 cells, which have a bursting phenotype but are largely distinct from the Chrna2 subset (Fig. 2), may belong to the Sst-MET 5 type. L4 X94 cells seem to correspond to Sst-MET type 8, based on their laminar position, dense axonal arbor in L4, and HPSE expression (Naka et al., 2019), although in visual cortex (used for the MET classification) they also have substantial axonal arborization in L1 (Scala et al., 2019). L5b X94 cells likely correspond to Sst-MET 7, based on L4-targeting and HPSE expression; but there are also HPSE-expressing, L4-targeting infragranular SOM cells in Sst-MET 11 and 12. The proposed correspondence of the 4 SOM subsets (including X98) to Sst-MET 1-8 is summarized in ***Table 7***. Notably, the descriptors “T-shaped” and “fanning out” do not map onto unique MET types; for example, Sst-MET 4, 6 and 7 contain “T-shaped” cells, while most Sst-MET 4 and 5, and some Sst-MET 9 and 12 cells are “fanning out” cells (Gouwens et al., 2020). Thus, we suggest that these terms be used as morphological descriptors and not as subtype designations.

**Table 7.** Suggested correspondence of SOM subtypes with the Sst-MET taxonomy. (Gouwens et al., 2020). Intersectional and transgenic subtypes are listed under the most *comparable* Sst-MET type; when listed in parenthesis, the correspondence is conjectural. Only Sst-MET types 1-8 are included.

|  |  |  |  |
| --- | --- | --- | --- |
| **MET 1**<br>Long-range | **MET 2**<br>(L2/3 Calb2) | **MET 3**<br>L2/3 Calb2 | **MET 4**<br>L5 Calb2 |
| **MET 5**<br>(X98) | **MET 6**<br>Chrna2 | **MET 7**<br>L5 X94 | **MET 8**<br>L4 X94 |

Our study is the first to apply retrograde labeling from an epipial dye deposit to identify L1-projecting interneurons in the mouse, although a previous study has done so in the rat (Ramos-Moreno and Clasca, 2014). L1-projecting SOM interneurons with radially ascending axons are historically referred to as Martinotti cells (Marin-Padilla, 1990; DeFelipe, 2002), and some earlier studies regarded “Martinotti” and “somatostatin-containing” as synonymous terms. That not all SOM interneurons are L1-projecting was made clear by the discovery of two non-L1-projecting SOM groups: the long-range-projecting, sleep-active, NPY/nNOS-expressing cells in L2 and L6 (Tomioka et al., 2005; Gerashchenko et al., 2008), and L4-projecting X94 cells in L4 and L5b (Ma et al., 2006). The recent transcriptomic studies revealed additional SOM types which do not project to L1. Out of the 13 Sst-MET types (Gouwens et al., 2020), 4 (Sst-MET 10-13) have cell bodies in L5/6 and axonal arborization concentrated in L4-L6, with virtually no axonal projections in L1. This is in addition to MET types 1 and 8 which have only minor projections in L1. Thus, only about half of all Sst-MET types are bona-fide Martinotti cells. The transcriptomic studies, however, are unable to estimate the relative abundance of different types, as cells in these studies were not sampled in an unbiased manner. Our retrograde labeling results provide, for the first time, an unbiased estimate of the fraction of mouse SOM interneurons with an axonal arbor in L1, which in our experiments was slightly over 50% and varied by layer, from 80% in L2/3 to 60% in L5, 40% in L6 and <20% in L4. Interestingly, the previous rat study (Ramos-Moreno and Clasca, 2014) only identified 26% of SOM cells in S1 as retrogradely labeled after an epipial FB deposit, most of them in L2/3, with none in L6. Both studies used the same dye concentration, and survival time in the rat study was 7 days, compared to 24 hours in the current study. It is possible that the thicker pia mater in the rat restricted diffusion of dye (and thereby dye uptake) in this previous study.

This study was conducted without prior knowledge of a comprehensive study of SOM subtypes recently posted in preprint form (Wu et al., 2022). Wu et al. used intersectional strategies to target 8 different transcriptomic SOM subtypes and to examine in detail synaptic inputs and outputs of 3 of them, including the Calb2 and Chrna2 subtypes examined here. Where overlapping, the results of our studies are in good agreement, except that we observed unequivocal separation in electrophysiological properties between the Calb2 and Chrna2 subtypes (***Figs. 7-9***) whereas Wu et al. did not. Both studies remain limited by existing driver lines, which may not be specific enough to target “pure” SOM subtypes. For example, the Calb2 subset contains both L2/3 and L5 neurons. While Wu et al. found that these subsets (which fall into different MET types) have very similar electrophysiological properties (Wu et al., 2022), they could still differ in their synaptic inputs and outputs. Indeed, it is suggested that L2/3 and L5 Martinotti cells preferentially target cells in layers 2/3 and 5, respectively (Jiang et al., 2015). Importantly, neither study has found a genetic strategy to specifically target the L4-targeting X94 subtype (Ma et al., 2006); while the Pdyn subset is enriched in X94-like neurons, both studies found that it includes many L1-projecting cells as well. Clearly, more specific driver lines and/or more sophisticated combinatorial reporters are needed to restrict reporter expression to smaller, more homogeneous populations. Once such genetic strategies are developed, they should be used to examine the local and long-range synaptic connections of the targeted populations, to document their pattern of activity in the behaving animal and to test their involvement in cortical computations related to sensory processing, motor planning and cognition. While a daunting task, accomplishing it is essential if we are to establish which of the different transcriptomic groups proposed by recent taxonomies are bona-fide, biologically meaningful neuronal subtypes, what roles each subtype plays in cortical computations and how it contributes to animal and human behavior.

## Supporting information

Extended data for Figure 6

Extended data for Figure 8

## ACKNOWLEDGEMENTS

We thank Qingyan Wang and Grace Jones for excellent technical support, Amanda Ammer for technical help with imaging, and Bernardo Martinez for assistance with graphics software. The Chrna2-cre animals were generously provided by J. Josh Lawrence. This study was supported by National Institutes of Health grant NS116604 to AA. Additional funding was provided by Transition Grant Support from the Office of Research and Graduate Education, WVU Health Sciences Center, and by the Program to Stimulate Competitive Research, Office of Research, WVU. REH was supported by NIH training grants GM081741 and GM132494. Confocal imaging was performed at the WVU Microscopic Imaging Facility, which was supported by NIH grants GM121322, GM103434, GM103503, GM104942, and RR016440.

## ETHICS STATEMENT

Animal husbandry and experimental procedures followed the Public Health Service Policy on Humane Care and Use of Laboratory Animals, and were approved by the WVU Institutional Animal Care and Use Committee. West Virginia University has a PHS-approved Animal Welfare Assurance D16-00362 (A3597-01).

## EXTENDED DATA LEGENDS

**Fig. 6-1. Summary data used to create the sunburst charts in Fig. 6**.

The table indicates the average number of cells counted in each layer for each antibody, for every combinations of genetic label (GFP+ or tdTomato+), retrograde label (FB+ or FB-) and immunostaining (Ab+ and Ab-), expressed as a fraction of all cells (tdTomato-expressing and GFP-expressing) counted in the same sections. N=3 for each antibody.

**Fig. 8-1. Summary of electrophysiological parameters measured in 94 neurons from the 4 genetic subsets.** See Methods for definition of the 10 parameters. In addition, the table lists the number of rebound spikes fired upon recovery from a hyperpolarizing current step; the results of applying the Decision Tree classifier in ***Fig. 9***, with wrong predictions highlighted in red; and the values of the largest two principal components and the largest two discriminant functions for each cell.

## REFERENCES

Abbas AI, Sundiang MJM, Henoch B, Morton MP, Bolkan SS, Park AJ, Harris AZ, Kellendonk C, Gordon JA (2018) Somatostatin Interneurons Facilitate Hippocampal-Prefrontal Synchrony and Prefrontal Spatial Encoding. Neuron 100:926–939 e923. doi:10.1016/j.neuron.2018.09.029

Adesnik H, Bruns W, Taniguchi H, Huang ZJ, Scanziani M (2012) A neural circuit for spatial summation in visual cortex. Nature 490:226–231. doi:10.1038/nature11526

Adler A, Zhao R, Shin ME, Yasuda R, Gan WB (2019) Somatostatin-Expressing Interneurons Enable and Maintain Learning-Dependent Sequential Activation of Pyramidal Neurons. Neuron. doi:10.1016/j.neuron.2019.01.036

Agmon A, Connors BW (1991) Thalamocortical responses of mouse somatosensory (barrel) cortex in vitro. Neuroscience 41:365–379.

Anderson KM, Collins MA, Kong R, Fang K, Li J, He T, Chekroud AM, Yeo BTT, Holmes AJ (2020) Convergent molecular, cellular, and cortical neuroimaging signatures of major depressive disorder. Proceedings of the National Academy of Sciences of the United States of America. doi:10.1073/pnas.2008004117

Artinian J, Jordan A, Khlaifia A, Honore E, La Fontaine A, Racine AS, Laplante I, Lacaille JC (2019) Regulation of Hippocampal Memory by mTORC1 in Somatostatin Interneurons. J Neurosci 39:8439–8456. doi:10.1523/JNEUROSCI.0728-19.2019

Asgarian Z, Oliveira MG, Stryjewska A, Maragkos I, Rubin AN, Magno L, Pachnis V, Ghorbani M, Hiebert SW, Denaxa M, Kessaris N (2022) MTG8 interacts with LHX6 to specify cortical interneuron subtype identity. Nature communications 13:5217. doi:10.1038/s41467-022-32898-6

Baimbridge KG, Celio MR, Rogers JH (1992) Calcium-binding proteins in the nervous system. Trends Neurosci 15:303–308.

Bloss EB, Cembrowski MS, Karsh B, Colonell J, Fetter RD, Spruston N (2016) Structured Dendritic Inhibition Supports Branch-Selective Integration in CA1 Pyramidal Cells. Neuron 89:1016–1030. doi:10.1016/j.neuron.2016.01.029

Cauller LJ, Clancy B, Connors BW (1998) Backward cortical projections to primary somatosensory cortex in rats extend long horizontal axons in layer I. J Comp Neurol 390:297–310.

Chen D, Wang C, Li M, She X, Yuan Y, Chen H, Zhang W, Zhao C (2019) Loss of Foxg1 Impairs the Development of Cortical SST-Interneurons Leading to Abnormal Emotional and Social Behaviors. Cereb Cortex 29:3666–3682. doi:10.1093/cercor/bhz114

Chen G, Zhang Y, Li X, Zhao X, Ye Q, Lin Y, Tao HW, Rasch MJ, Zhang X (2017) Distinct Inhibitory Circuits Orchestrate Cortical beta and gamma Band Oscillations. Neuron 96:1403–1418 e1406. doi:10.1016/j.neuron.2017.11.033

Cummings KA, Clem RL (2020) Prefrontal somatostatin interneurons encode fear memory. Nat Neurosci 23:61–74. doi:10.1038/s41593-019-0552-7

Daigle TL et al. (2018) A Suite of Transgenic Driver and Reporter Mouse Lines with Enhanced Brain-Cell-Type Targeting and Functionality. Cell 174:465–480 e422. doi:10.1016/j.cell.2018.06.035

DeFelipe J (2002) Cortical interneurons: from Cajal to 2001. Prog Brain Res 136:215–238.

Demeulemeester H, Vandesande F, Orban GA, Brandon C, Vanderhaeghen JJ (1988) Heterogeneity of GABAergic cells in cat visual cortex. J Neurosci 8:988-1000.

Dobrzanski G, Lukomska A, Zakrzewska R, Posluszny A, Kanigowski D, Urban-Ciecko J, Liguz-Lecznar M, Kossut M (2022) Learning-induced plasticity in the barrel cortex is disrupted by inhibition of layer 4 somatostatin-containing interneurons. Biochim Biophys Acta Mol Cell Res 1869:119146. doi:10.1016/j.bbamcr.2021.119146

Doron M, Chindemi G, Muller E, Markram H, Segev I (2017) Timed Synaptic Inhibition Shapes NMDA Spikes, Influencing Local Dendritic Processing and Global I/O Properties of Cortical Neurons. Cell reports 21:1550–1561. doi:10.1016/j.celrep.2017.10.035

Druckmann S, Hill S, Schurmann F, Markram H, Segev I (2013) A hierarchical structure of cortical interneuron electrical diversity revealed by automated statistical analysis. Cereb Cortex 23:2994–3006. doi:10.1093/cercor/bhs290

Fee C, Banasr M, Sibille E (2017) Somatostatin-Positive Gamma-Aminobutyric Acid Interneuron Deficits in Depression: Cortical Microcircuit and Therapeutic Perspectives. Biological psychiatry. doi:10.1016/j.biopsych.2017.05.024

Freund TF, Buzsaki G (1996) Interneurons of the hippocampus. Hippocampus 6:347–470. doi:10.1002/(SICI)1098-1063(1996)6:4<347::AID-HIPO1>3.0.CO;2-I

Gerashchenko D, Wisor JP, Burns D, Reh RK, Shiromani PJ, Sakurai T, de la Iglesia HO, Kilduff TS (2008) Identification of a population of sleep-active cerebral cortex neurons. Proceedings of the National Academy of Sciences of the United States of America 105:10227–10232. doi:10.1073/pnas.0803125105

Gonchar Y, Wang Q, Burkhalter A (2007) Multiple distinct subtypes of GABAergic neurons in mouse visual cortex identified by triple immunostaining. Frontiers in neuroanatomy 1:3. doi:10.3389/neuro.05.003.2007

Good PI (1999) Resampling Methods: A Practical Guide to Data Analysis. Boston: Birkhauser.

Gouwens NW et al. (2020) Integrated Morphoelectric and Transcriptomic Classification of Cortical GABAergic Cells. Cell 183:935–953 e919. doi:10.1016/j.cell.2020.09.057

Hakim R, Shamardani K, Adesnik H (2018) A neural circuit for gamma-band coherence across the retinotopic map in mouse visual cortex. eLife 7. doi:10.7554/eLife.28569

Halabisky B, Shen F, Huguenard JR, Prince DA (2006) Electrophysiological classification of somatostatin-positive interneurons in mouse sensorimotor cortex. J Neurophysiol 96:834–845. doi:10.1152/jn.01079.2005

He L, Caudill MS, Jing J, Wang W, Sun Y, Tang J, Jiang X, Zoghbi HY (2022) A weakened recurrent circuit in the hippocampus of Rett syndrome mice disrupts long-term memory representations. Neuron 110:1689–1699 e1686. doi:10.1016/j.neuron.2022.02.014

He M, Tucciarone J, Lee S, Nigro MJ, Kim Y, Levine JM, Kelly SM, Krugikov I, Wu P, Chen Y, Gong L, Hou Y, Osten P, Rudy B, Huang ZJ (2016) Strategies and Tools for Combinatorial Targeting of GABAergic Neurons in Mouse Cerebral Cortex. Neuron 91:1228–1243. doi:10.1016/j.neuron.2016.08.021

Hilscher MM, Leao RN, Edwards SJ, Leao KE, Kullander K (2017) Chrna2-Martinotti Cells Synchronize Layer 5 Type A Pyramidal Cells via Rebound Excitation. PLoS Biol 15:e2001392. doi:10.1371/journal.pbio.2001392

Hu H, Cavendish JZ, Agmon A (2013) Not all that glitters is gold: off-target recombination in the somatostatin-IRES-Cre mouse line labels a subset of fast-spiking interneurons. Frontiers in neural circuits 7:195. doi:10.3389/fncir.2013.00195

Isaacson JS, Scanziani M (2011) How inhibition shapes cortical activity. Neuron 72:231–243. doi:10.1016/j.neuron.2011.09.027

Jadi M, Polsky A, Schiller J, Mel BW (2012) Location-dependent effects of inhibition on local spiking in pyramidal neuron dendrites. PLoS Comput Biol 8:e1002550. doi:10.1371/journal.pcbi.1002550

Jiang X, Shen S, Cadwell CR, Berens P, Sinz F, Ecker AS, Patel S, Tolias AS (2015) Principles of connectivity among morphologically defined cell types in adult neocortex. Science 350:aac9462. doi:10.1126/science.aac9462

Kato HK, Asinof SK, Isaacson JS (2017) Network-Level Control of Frequency Tuning in Auditory Cortex. Neuron 95:412–423 e414. doi:10.1016/j.neuron.2017.06.019

Kawaguchi Y, Kubota Y (1996) Physiological and morphological identification of somatostatin- or vasoactive intestinal polypeptide-containing cells among GABAergic cell subtypes in rat frontal cortex. J Neurosci 16:2701–2715.

Kawaguchi Y, Kubota Y (1997) GABAergic cell subtypes and their synaptic connections in rat frontal cortex. Cereb Cortex 7:476–486.

Kepecs A, Fishell G (2014) Interneuron cell types are fit to function. Nature 505:318–326. doi:10.1038/nature12983

Krashes MJ, Shah BP, Madara JC, Olson DP, Strochlic DE, Garfield AS, Vong L, Pei H, Watabe-Uchida M, Uchida N, Liberles SD, Lowell BB (2014) An excitatory paraventricular nucleus to AgRP neuron circuit that drives hunger. Nature 507:238–242. doi:10.1038/nature12956

Kubota Y, Hattori R, Yui Y (1994) Three distinct subpopulations of GABAergic neurons in rat frontal agranular cortex. Brain Res 649:159–173.

Lakunina AA, Nardoci MB, Ahmadian Y, Jaramillo S (2020) Somatostatin-expressing interneurons in the auditory cortex mediate sustained suppression by spectral surround. J Neurosci. doi:10.1523/JNEUROSCI.1735-19.2020

Le Magueresse C, Monyer H (2013) GABAergic interneurons shape the functional maturation of the cortex. Neuron 77:388–405. doi:10.1016/j.neuron.2013.01.011

Leao RN, Mikulovic S, Leao KE, Munguba H, Gezelius H, Enjin A, Patra K, Eriksson A, Loew LM, Tort AB, Kullander K (2012) OLM interneurons differentially modulate CA3 and entorhinal inputs to hippocampal CA1 neurons. Nat Neurosci 15:1524–1530. doi:10.1038/nn.3235

Lovett-Barron M, Turi GF, Kaifosh P, Lee PH, Bolze F, Sun XH, Nicoud JF, Zemelman BV, Sternson SM, Losonczy A (2012) Regulation of neuronal input transformations by tunable dendritic inhibition. Nat Neurosci 15:423–430, S421-423. doi:10.1038/nn.3024

Ma Y, Hu H, Berrebi AS, Mathers PH, Agmon A (2006) Distinct subtypes of somatostatin-containing neocortical interneurons revealed in transgenic mice. J Neurosci 26:5069–5082. doi:10.1523/JNEUROSCI.0661-06.2006

Machold R, Dellal S, Valero M, Zurita H, Kruglikov I, Meng J, Hanson JL, Hashikawa Y, Schuman B, Buzsáki G, Rudy B (2022) Id2 GABAergic interneurons: a neglected fourth major group of cortical inhibitory cells. bioRxiv:2022.2012.2001.518752. doi:10.1101/2022.12.01.518752

Madisen L, Zwingman TA, Sunkin SM, Oh SW, Zariwala HA, Gu H, Ng LL, Palmiter RD, Hawrylycz MJ, Jones AR, Lein ES, Zeng H (2010) A robust and high-throughput Cre reporting and characterization system for the whole mouse brain. Nat Neurosci 13:133–140. doi:10.1038/nn.2467

Makino H, Komiyama T (2015) Learning enhances the relative impact of top-down processing in the visual cortex. Nat Neurosci 18:1116–1122. doi:10.1038/nn.4061

Manly BFJ (2005) Multivariate Statistical Methods: A Primer. Boca Raton: Chapman &Hall/CRC

Marin-Padilla M (1990) The pyramidal cell and its local-circuit interneurons: a hypothetical unit of the Mammalian cerebral cortex. J Cogn Neurosci 2:180–194. doi:10.1162/jocn.1990.2.3.180

McGarry LM, Packer AM, Fino E, Nikolenko V, Sippy T, Yuste R (2010) Quantitative classification of somatostatin-positive neocortical interneurons identifies three interneuron subtypes. Frontiers in neural circuits 4:12. doi:10.3389/fncir.2010.00012

Mikulovic S, Restrepo CE, Hilscher MM, Kullander K, Leao RN (2015) Novel markers for OLM interneurons in the hippocampus. Frontiers in cellular neuroscience 9:201. doi:10.3389/fncel.2015.00201

Mitchell BD, Cauller LJ (2001) Corticocortical and thalamocortical projections to layer I of the frontal neocortex in rats. Brain Res 921:68–77.

Muller-Komorowska D, Opitz T, Elzoheiry S, Schweizer M, Ambrad Giovannetti E, Beck H (2020) Nonspecific Expression in Limited Excitatory Cell Populations in Interneuron-Targeting Cre-driver Lines Can Have Large Functional Effects. Frontiers in neural circuits 14:16. doi:10.3389/fncir.2020.00016

Munoz W, Tremblay R, Levenstein D, Rudy B (2017) Layer-specific modulation of neocortical dendritic inhibition during active wakefulness. Science 355:954–959. doi:10.1126/science.aag2599

Naka A, Veit J, Shababo B, Chance RK, Risso D, Stafford D, Snyder B, Egladyous A, Chu D, Sridharan S, Mossing DP, Paninski L, Ngai J, Adesnik H (2019) Complementary networks of cortical somatostatin interneurons enforce layer specific control. eLife 8. doi:10.7554/eLife.43696

Nigro MJ, Hashikawa-Yamasaki Y, Rudy B (2018) Diversity and Connectivity of Layer 5 Somatostatin-Expressing Interneurons in the Mouse Barrel Cortex. J Neurosci 38:1622–1633. doi:10.1523/JNEUROSCI.2415-17.2017

Oberlaender M, de Kock CP, Bruno RM, Ramirez A, Meyer HS, Dercksen VJ, Helmstaedter M, Sakmann B (2012) Cell type-specific three-dimensional structure of thalamocortical circuits in a column of rat vibrissal cortex. Cereb Cortex 22:2375–2391. doi:10.1093/cercor/bhr317

Oliva AA, Jr., Jiang M, Lam T, Smith KL, Swann JW (2000) Novel hippocampal interneuronal subtypes identified using transgenic mice that express green fluorescent protein in GABAergic interneurons. J Neurosci 20:3354–3368.

Paluszkiewicz SM, Olmos-Serrano JL, Corbin JG, Huntsman MM (2011) Impaired inhibitory control of cortical synchronization in fragile X syndrome. J Neurophysiol 106:2264–2272. doi:10.1152/jn.00421.2011

Plummer NW, Evsyukova IY, Robertson SD, de Marchena J, Tucker CJ, Jensen P (2015) Expanding the power of recombinase-based labeling to uncover cellular diversity. Development 142:4385–4393. doi:10.1242/dev.129981

Porter JT, Johnson CK, Agmon A (2001) Diverse types of interneurons generate thalamus-evoked feedforward inhibition in the mouse barrel cortex. J Neurosci 21:2699–2710.

Ramos-Moreno T, Clasca F (2014) Quantitative mapping of the local and extrinsic sources of GABA and Reelin to the layer Ia neuropil in the adult rat neocortex. Brain structure & function 219:1639–1657. doi:10.1007/s00429-013-0591-x

Reep RL (2000) Cortical layer VII and persistent subplate cells in mammalian brains. Brain Behav Evol 56:212–234. doi:10.1159/000047206

Riedemann T, Schmitz C, Sutor B (2016) Immunocytochemical heterogeneity of somatostatin-expressing GABAergic interneurons in layers II and III of the mouse cingulate cortex: A combined immunofluorescence/design-based stereologic study. J Comp Neurol 524:2281–2299. doi:10.1002/cne.23948

Riedemann T, Straub T, Sutor B (2018) Two types of somatostatin-expressing GABAergic interneurons in the superficial layers of the mouse cingulate cortex. PloS one 13:e0200567. doi:10.1371/journal.pone.0200567

Rudy B, Fishell G, Lee S, Hjerling-Leffler J (2011) Three groups of interneurons account for nearly 100% of neocortical GABAergic neurons. Developmental neurobiology 71:45–61. doi:10.1002/dneu.20853

Scala F, Kobak D, Shan S, Bernaerts Y, Laturnus S, Cadwell CR, Hartmanis L, Froudarakis E, Castro JR, Tan ZH, Papadopoulos S, Patel SS, Sandberg R, Berens P, Jiang X, Tolias AS (2019) Layer 4 of mouse neocortex differs in cell types and circuit organization between sensory areas. Nature communications 10:4174. doi:10.1038/s41467-019-12058-z

Schuman B, Machold RP, Hashikawa Y, Fuzik J, Fishell GJ, Rudy B (2019) Four Unique Interneuron Populations Reside in Neocortical Layer 1. J Neurosci 39:125–139. doi:10.1523/JNEUROSCI.1613-18.2018

Sohn J, Hioki H, Okamoto S, Kaneko T (2014) Preprodynorphin-expressing neurons constitute a large subgroup of somatostatin-expressing GABAergic interneurons in the mouse neocortex. J Comp Neurol 522:1506–1526. doi:10.1002/cne.23477

Taniguchi H, He M, Wu P, Kim S, Paik R, Sugino K, Kvitsiani D, Fu Y, Lu J, Lin Y, Miyoshi G, Shima Y, Fishell G, Nelson SB, Huang ZJ (2011) A resource of Cre driver lines for genetic targeting of GABAergic neurons in cerebral cortex. Neuron 71:995–1013. doi:10.1016/j.neuron.2011.07.026

Tasic B et al. (2018) Shared and distinct transcriptomic cell types across neocortical areas. Nature 563:72–78. doi:10.1038/s41586-018-0654-5

Thomson AM (2010) Neocortical layer 6, a review. Frontiers in neuroanatomy 4:13. doi:10.3389/fnana.2010.00013

Tomioka R, Okamoto K, Furuta T, Fujiyama F, Iwasato T, Yanagawa Y, Obata K, Kaneko T, Tamamaki N (2005) Demonstration of long-range GABAergic connections distributed throughout the mouse neocortex. Eur J Neurosci 21:1587–1600. doi:10.1111/j.1460-9568.2005.03989.x

Tremblay R, Lee S, Rudy B (2016) GABAergic Interneurons in the Neocortex: From Cellular Properties to Circuits. Neuron 91:260–292. doi:10.1016/j.neuron.2016.06.033

van Brederode JF, Helliesen MK, Hendrickson AE (1991) Distribution of the calcium-binding proteins parvalbumin and calbindin-D28k in the sensorimotor cortex of the rat. Neuroscience 44:157–171.

Veit J, Hakim R, Jadi MP, Sejnowski TJ, Adesnik H (2017) Cortical gamma band synchronization through somatostatin interneurons. Nat Neurosci. doi:10.1038/nn.4562

Wang Y, Ye M, Kuang X, Li Y, Hu S (2018) A simplified morphological classification scheme for pyramidal cells in six layers of primary somatosensory cortex of juvenile rats. IBRO Rep 5:74–90. doi:10.1016/j.ibror.2018.10.001

Wang Y, Toledo-Rodriguez M, Gupta A, Wu C, Silberberg G, Luo J, Markram H (2004) Anatomical, physiological and molecular properties of Martinotti cells in the somatosensory cortex of the juvenile rat. The Journal of physiology 561:65–90. doi:10.1113/jphysiol.2004.073353

Wengert ER, Miralles RM, Wedgwood KCA, Wagley PK, Strohm SM, Panchal PS, Idrissi AM, Wenker IC, Thompson JA, Gaykema RP, Patel MK (2021) Somatostatin-Positive Interneurons Contribute to Seizures in SCN8A Epileptic Encephalopathy. J Neurosci 41:9257–9273. doi:10.1523/JNEUROSCI.0718-21.2021

Wu SJ et al. (2022) Cortical somatostatin interneuron subtypes form cell-type specific circuits. bioRxiv:2022.2009.2029.510081. doi:10.1101/2022.09.29.510081

Xu X, Roby KD, Callaway EM (2006) Mouse cortical inhibitory neuron type that coexpresses somatostatin and calretinin. J Comp Neurol 499:144–160. doi:10.1002/cne.21101

Xu X, Roby KD, Callaway EM (2010) Immunochemical characterization of inhibitory mouse cortical neurons: three chemically distinct classes of inhibitory cells. J Comp Neurol 518:389–404. doi:10.1002/cne.22229

Zorrilla de San Martin J, Donato C, Peixoto J, Aguirre A, Choudhary V, De Stasi AM, Lourenco J, Potier MC, Bacci A (2020) Alterations of specific cortical GABAergic circuits underlie abnormal network activity in a mouse model of Down syndrome. eLife 9. doi:10.7554/eLife.58731

